# Microfluidic-Induced Sleep: A Spontaneous *C. elegans* Sleep State Regulated by Satiety, Thermosensation and Mechanosensation

**DOI:** 10.1101/547075

**Authors:** Daniel L. Gonzales, Jasmine Zhou, Bo Fan, Jacob T. Robinson

## Abstract

One remarkable feature of behavior is an animal’s ability to rapidly switch between activity states; however, how the brain regulates these spontaneous transitions based on the animal’s perceived environment is not well understood. Here we show a *C. elegans* sleep-like state on a scalable platform that enables simultaneous control of multiple environmental factors including temperature, mechanical stress, and food availability. This brief quiescent state, we refer to as “microfluidic-induced sleep,” occurs spontaneously in microfluidic chambers, which allows us to track animal movement and perform whole-brain imaging. With these capabilities, we establish that microfluidic-induced sleep meets the behavioral requirements of *C. elegans* sleep and depends on multiple factors, such as satiety and temperature. Additionally, we show for the first time that *C. elegans* sleep can be induced through mechanosensory pathways. Together, these results establish a rich model system for studying how animals process multiple sensory pathways to regulate behavioral states.

## Introduction

Understanding how animals select behaviors based on sensory information is a fundamental goal of neuroscience ^1–4;^ however, sensorimotor transformations can vary dramatically depending on the state of the animal’s nervous system ^5–8^. Wakefulness and arousal ^9, 10^, locomotor activity states ^11, 12^, satiety ^13^, attention ^14^, and emotions ^5, 6, 15^ represent a spectrum of physiological and neural states that can dramatically affect how animals respond to a given stimuli. Small animals like the nematode *C. elegans* are tractable model organisms for understanding how physiological and neural states combine with information from multiple sensory pathways and give rise to specific behavior ^3, 6, 8, 13, 16–21^.

Sleep is one example of a neural state that dramatically alters an animal’s sensorimotor transformations ^22, 23^. Studies of sleep across the phylogenetic tree have shown that sensory systems transition to a reduced activity state that leads to a decreased animal response to external stimuli ^24–28^. Additionally, sleep and wakefulness have been shown to correspond with distinct patterns of neural activity in humans ^22, 23^, cats ^29^, rodents ^23^, and fruit flies ^30, 31^. Likewise, whole-brain recordings from *C. elegans* suggest that the majority of worm brain activity can be represented on a low-dimensional manifold ^32^, and that the activity on this manifold shifts from phasic to fixed-point attractor dynamics during developmental sleep ^33^. Additionally, studies with *C. elegans* have revealed molecular pathways ^25, 34–37^, neural circuits ^34, 38–43^, and neuropeptides ^39, 42, 44, 45^ that drive nematode sleep and arousal, and some of these mechanisms are conserved in in other animals ^46–48^. These reports have paved the way for using *C. elegans* sleep as a model system to understand spontaneous brain-state transitions between sleep and wakefulness. Given the number of unique regulators of *C. elegans* sleep, there is a need to identify a behavior that facilitates our understanding of how brain-wide neural circuits transduce multiple external and internal factors to drive sleep-wake transitions. Here, we describe a *C. elegans* sleep behavior in a microfluidic environment, termed “microfluidic-induced sleep,” that is amenable to whole-brain imaging and is regulated both by the animal’s physiological state and external sensory queues. While microfluidic-induced sleep is likely a form of previously-reported nematode quiescence behaviors, such as stress-induced sleep and episodic swimming ^40, 49^, the rapid rate of state transitions and the fact that these quiescence behaviors occur in microfluidic devices provides and number of advantages for studying the mechanisms of state transitions with precise environmental controllability, high-throughput screenings and whole-brain imaging.

## Results

When confined to microfluidic chambers we found that adult *C. elegans* rapidly and spontaneously switch between normal activity and brief quiescent bouts without any additional stimuli (**Figure 1A**). We initially observed this behavior when immobilizing worms for recording body-wall muscle electrophysiology ^50^. In these electrical recordings we observed minutes-long periods of muscle inactivity that corresponded to whole-animal quiescence ^50^. We have since found that this quiescence occurs in a wide range of microfluidic geometries (**Supplementary Movie 1-Supplementary Movie 2**).

**Figure 1.**
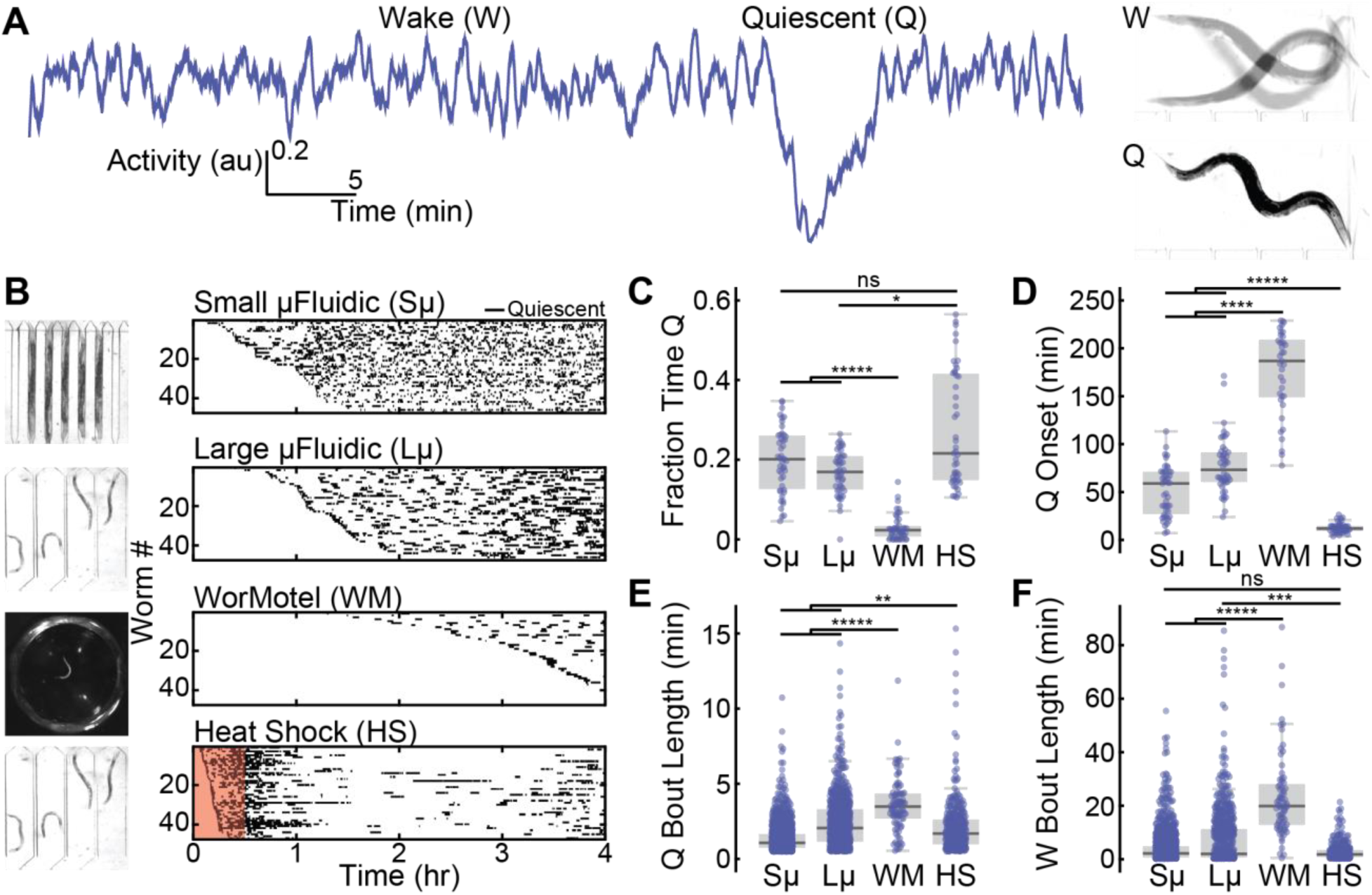
Quiescence dynamics are strongly affected by microfluidic environments. *(A)* Characteristic worm quiescence in a microfluidic chamber. *(Left)* 1 hr activity trace from a worm swimming in a microfluidic chamber, calculated by subtracting consecutive frames to quantify movement (see Methods). Quiescence is hallmarked by a clear drop in animal activity to near zero. *(Right)* 1 s overlap of frames shows the animal swimming during the wake period (W) and immobile during quiescence (Q). *(B)* All quiescent bouts recorded from animals in a small microfluidic chamber (50 μm width), large microfluidic chamber (500 μm width), WorMotel multi-well PDMS device, and large microfluidic device during a 30 min heat shock at 30 °C (see inset images). Raster plots show the bouts recorded from each animal during a 4 hr imaging period (n = 47 animals for each condition). Experiments represented in the raster plots are sorted by the onset of the first detected sleep bout. *(C-F)* Quantitative sleep metrics. Sμ = small microfluidic chambers, Lμ = large microfluidic chambers, WM = WorMotel, HS = first hour of heat shock. *(C)* The total fraction of time each animal spent in the quiescent state. *(D)* Onset time of the first quiescent bout. *(E)* Length of individual quiescent bouts. *(F)* Length of individual wake bouts, excluding the first period of wake after animals are loaded into the chamber. (n = 47 animals for each condition; *p < 0.05, ** p < 0.01, ***p<0.001, ****p<0.0001, *****p<0.00001; Kruskal-Wallis with a *post-hoc* Dunn-Sidak test).

We found that the onset, frequency, and duration of quiescent bouts in microfluidic chambers are unique compared to previously-reported sleep behaviors in adult *C. elegans* (**Figure 1**). While larval *C. elegans* display quiescence during lethargus ^25^, quiescence in adult worms occurs in only a few situations, such as after several hours of swimming ^49, 51–53^ or after exposure to extreme environmental conditions ^40, 44, 45, 54, 55^. To determine how microfluidic-induced quiescence compares to the well-established “episodic behavior” that occurs in swimming animals and “stress-induced quiescence” that occurs from a 30 min heat shock, we quantified these behaviors using the same methods to analyze microfluidic-induced sleep (**Figure 1B-F**). During a 4 hr imaging period, we found that animals partially immobilized in microfluidic channels (50 μm width, ∼0.004 μL per chamber, **Supplementary Movie 2**) displayed short (1.3 ± 0.1 min, mean ± sem, **Figure 1E**) bouts that, on average, began within the first hour of imaging (51 ± 4 min, mean ± sem, **Figure 1D**). Likewise, we also observed frequent, but longer (2.4 ± 0.3 min, mean ± sem, **Figure 1E**) quiescent bouts when the microfluidic chambers were large enough to allow animals to swim (500 μm width, ∼0.1 μL per chamber, **Supplementary Movie 1**). We compared these data to swimming-induced quiescence, where animals alternate between swimming and quiescence in a large multi-well device or “WorMotel” (**Figure 1B**, 8 μL of buffer per well) ^56^. In WorMotel, we recorded 5-7 times less sleep over the imaging period compared to microfluidic chambers (**Figure 1C**) and longer quiescent bouts compared to either microfluidic geometry (3.6 ± 0.3 min, mean ± sem, **Figure 1E**). Additionally, the average quiescence onset time was more than double what we observed for microfluidic-induced sleep (175 ± 6 min, mean ± sem, **Figure 1D**). For our final comparison point, we heat-shocked animals in the large microfluidic device for 30 min at 30 °C to induce stress-quiescence, then analyzed animal behavior during the heat shock plus a 30 min recovery period ^40^ (**Figure 1B**). As expected, during the noxious heat stimulus animals displayed their first quiescent bout more than 4-6 times faster than in microfluidic chambers with no external heat applied (13 ± 0.7 min, mean ± sem, **Figure 1D**).

These initial data show that compared to episodic swimming in WorMotel, microfluidic-induced sleep occurs with a higher frequency, faster onset and in shorter bouts (**Figure 1C-E**). Similarly, microfluidic-induced sleep onset occurs on a significantly different timescale compared to stress-induced sleep (**Figure 1B, D**). It is important to note that while the microfluidic-induced sleep phenotype shows a number of quantitative differences when compared to episodic swimming and stress-induced quiescence, it is not clear if microfluidic-induced quiescence can be classified as a new *C. elegans* behavior. It is possible that the microfluidic environment accelerates episodic swimming or introduces mild stressors that more slowly lead to stress-induced quiescence. Thus, we consider microfluidic-induced quiescence to merely display quantifiably distinct dynamics compared to other reported quiescent behaviors in *C. elegans* adults and is not necessarily a new behavioral state.

Our observation of spontaneous *C. elegans* quiescence in microfluidic chambers led us to determine whether this behavioral state transition meets the evolutionarily-conserved criteria for classification as sleep: reversibility, a decreased response to stimuli, homeostasis, and a stereotypical posture ^24^. *C. elegans* developmentally-timed sleep as larvae meets all requirements ^25, 26, 46, 57–62^. However, reports of sleep in adult worms have only observed reversibility and a decreased response to stimuli ^40, 46^. Here, we tested for a stereotyped posture, reversibility, decreased response to stimuli, and a homeostatic rebound.

Similar to previous reports ^57, 59, 60^, we found that animals exhibited a stereotyped posture during quiescence (**Figure 2A**). The body curvature of animals crawling on agar typically decreases during developmental sleep ^57^. In our case, we found that animals swimming in microfluidics show increased body curvature during sleep (from 3.9 ± 0.04 radians during wakefulness to 4.5 ± 0.1 radians during quiescence (mean ± sem), p < 0.0001, **Figure 2A**, right). To test for reversibility, we used blue light (5 s pulse, 5 mW/cm^2^) as a strong stimulus and found that this rapidly and reliably reversed the quiescent state, leading to a dramatic increase in both behavioral activity and nose speed (**Figure 2B**, **Supplementary Movie 3**). To test for a decreased response to stimuli, we fabricated microfluidic push-down valves designed to deliver tunable mechanical stimuli, similar to previously developed technology for worms ^63^. We found that when we applied a strong stimulus (high pressure, 30 psi), both wake and quiescent animals robustly responded with a significant increase in behavioral activity, which provides additional confirmation that the quiescent state is reversible (**Figure 2B**, **Supplementary Movie 4**). When we applied a weak stimulus to wake animals (low pressure, 15 psi), we again reliably recorded a significant increase in activity that matched the response of the strong stimulation (**Figure 2C**, **Supplementary Movie 5**). However, when we delivered a weak stimulus to quiescent animals, they responded weakly, exhibiting an average behavioral activity less than half that of animals in the awake state (**Figure 2C-D**, **Supplementary Movie 5**). In fact, less than 40% of quiescent animals that received a weak stimulus transitioned to wakefulness, compared to >75% of animals being in the awake state following stimulation for all other experimental conditions (**Figure 2E**). Importantly, because the strong stimuli evoked a similar strong behavioral response from both quiescent animals and awake animals, animals in the quiescent state are not less capable of responding to stimuli provided the stimuli is sufficiently intense (**Figure 2C-D**). Further analyses also showed that animals in the sleep state are less likely to respond to weak stimuli and transition to wakefulness, compared to animals in the wake state regardless of their activity level prior to the stimulus (**Supplementary Figure 1**). When we analyzed data from the wake animals, we found that nearly 70% of animals with activity levels below our sleep threshold for 3 seconds prior to the weak stimuli transitioned to wakefulness, compared to <40% for animals that we classified to be in the sleep state prior to a weak stimulus (**Supplementary Figure 1**). These results are consistent with quiescence being a sleep state rather than simply a low-activity state of wakefulness ^25, 26, 40, 61^.

**Figure 2.**
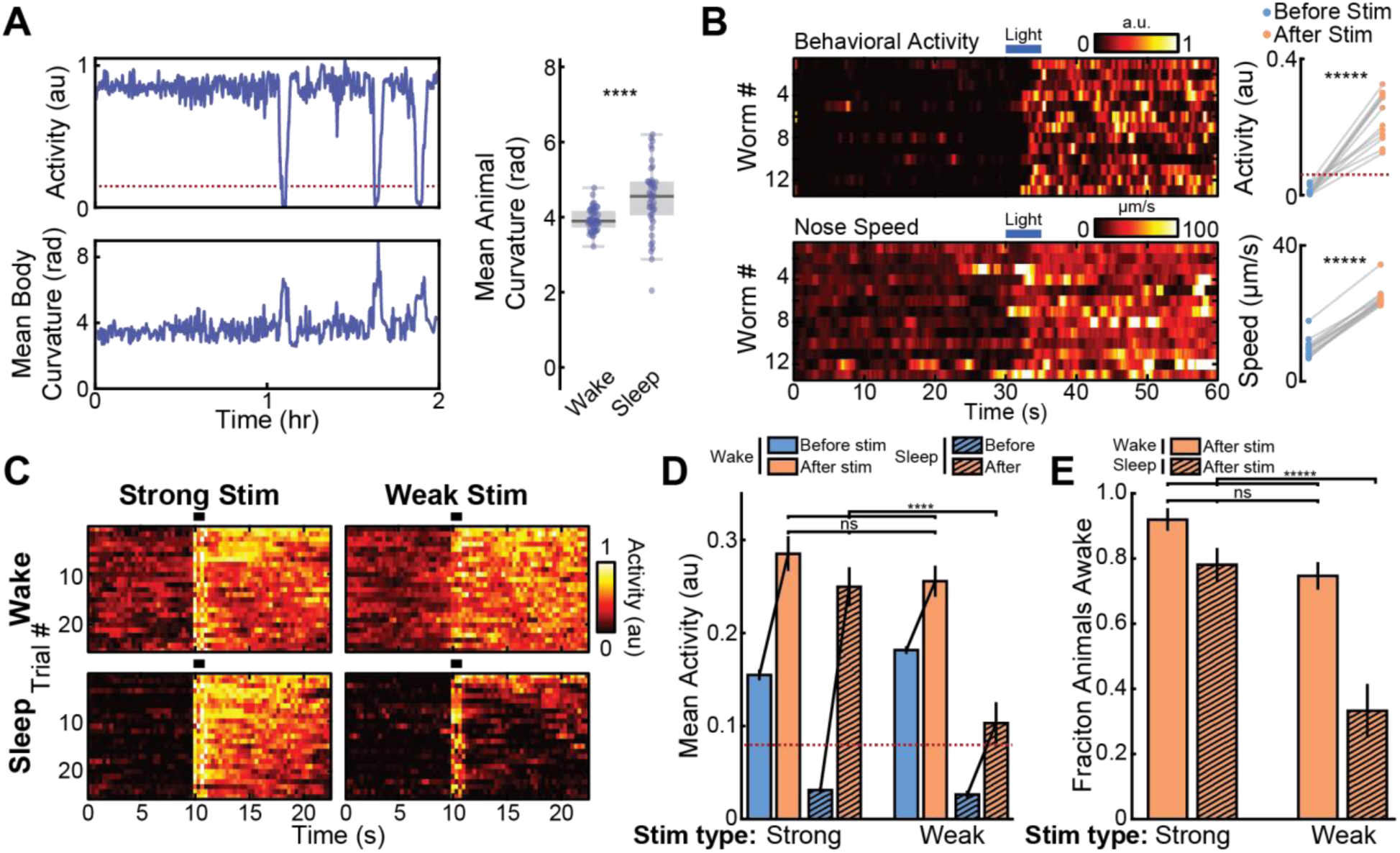
During microfluidic-induced sleep animals display stereotypical posture, reversibility, and a decreased response to weak stimuli. *(A) C. elegans* show increased body curvature during sleep. *(Left)* Representative example of quantified behavioral activity and normalized body curvature (see Methods) for a single animal during a 2 hr recording. Dotted line indicates sleep threshold. Increased body curvature correlates with sleep bouts. *(Right)* Across a population of animals, average body curvature increases during sleep (from “Large μFluidic” data set in Figure 1, n = 47 animals, ****p<0.0001, unpaired t-test). *(B)* The sleep state is reversible. *(Left)* Heatmaps of animal behavioral activity (top) and nose speed (bottom). Measurements begin with all animals in the quiescent state. The 5 s light pulse at t = 30 s results in a rapid increase in animal activity and nose speed. *(Right)* Mean behavioral activity (top) and nose speed (bottom) of each animal during the first and second 30 s of imaging shows a clear transition from sleep to wake states. Dotted line indicated sleep threshold. (n = 13 worms, *****p < 0.00001, paired t-test). *(C-E)* Quiescent animals have a decreased response to weak sensory stimuli. *(C)* Heatmaps of activity from awake and sleeping animals that received strong or weak mechanical stimuli from microfluidic valves (**Supplementary Movie 4-Supplementary Movie 5**). For each condition, only the 25 trials with the highest mean activity post-stimulation are shown. “Wake” indicates trials in which animals were awake but have a below-average activity before stimulation (see Methods). “Sleep” indicates trials in which animals were quiescent before stimulation. Heatmaps contain an ∼2 s long stimulation artifact beginning at 10 s due to movement from microfluidic valves. *(D)* Mean behavioral activity before and after mechanical stimulation from the trials in (C). Dotted line indicates sleep threshold. In all cases, average activity significantly increases after the stimulation (largest p-value = 0.002, paired two-sided t-test). However, following the stimulation behavioral activity is not significantly different in all cases other than when quiescent animals received weak mechanical stimuli (Error bars are sem; Strong-Wake n = 108, Strong-Sleep n = 51, Weak-Wake n = 176, Weak-Sleep n = 29; ****p < 0.0001, ns = not significant, Kruskal-Wallis with a *post-hoc* Dunn-Sidak test). *(E)* Fraction of animals in the wake state following mechanical stimuli. A significant difference only occurs in the case of quiescent animals receiving a weak mechanical stimuli. (Error bars are standard deviation, calculated by bootstrapping each data set with 5000 iterations; ns = not significant, *****p<0.00001; significance was calculated by data resampling 5000 iterations and a *post hoc* Bonferroni correction).

In addition, we also tested for homeostatic rebound, which is found in worm developmental sleep ^25, 43, 57^. Similar to previous studies with *C. elegans* developmental sleep ^57^, we hypothesized that longer wake bouts would lead to longer sleep bouts due to micro-homeostatic mechanisms. Indeed, we found that sleep bouts increased from 1.5 ± 0.1 min to 2.5 ± 0.1 min (mean ± sem) as the preceding wake bout increased in length from < 1 min to 20 min (**Supplementary Figure 2A**). However, even when wake bout lengths increased to longer than an hour, the sleep bouts lengths plateaued to an average of only 2.3 ± 0.1 min, contradicting the hypothesis that extended wake periods lead to sleep deprivation and homeostatic rebound. To directly test the effects of sleep deprivation, we performed optogenetic inhibition of the RIS neuron (**Supplementary Figure 2B-D**), which is known to be implicated in *C. elegans* sleep ^38, 39, 43^ (see also **Figure 3** and **Figure 5**). This optogenetic approach allowed us to have a control group (raised in the absence of all-trans retinal) that is also exposed to light but does not have optogenetic inhibition of the RIS neuron, thus controlling for any potential effects of stress produced by blue light illumination. We found that although RIS inhibition for 30 min did not fully abolish sleep, optically inhibited animals exhibited 324% more sleep and 230% more low-activity behavior compared to control animals during the refractory period following the optogenetic inhibition of the RIS neuron (p < 0.01) (**Supplementary Figure 2D**). RIS-inhibited animals also showed sleep bouts lasting 1.5 times longer than control animals (p < 0.01) (**Supplementary Figure 2C-D**). From our results combined, we conclude that animals show a homeostatic rebound in response to prolonged RIS inhibition. Therefore, microfluidic-induced quiescence indeed meets the behavioral precedents to be called a *C. elegans* sleep state: stereotyped posture, reversibility, a decreased response to stimuli, and homeostasis.

**Figure 3.**
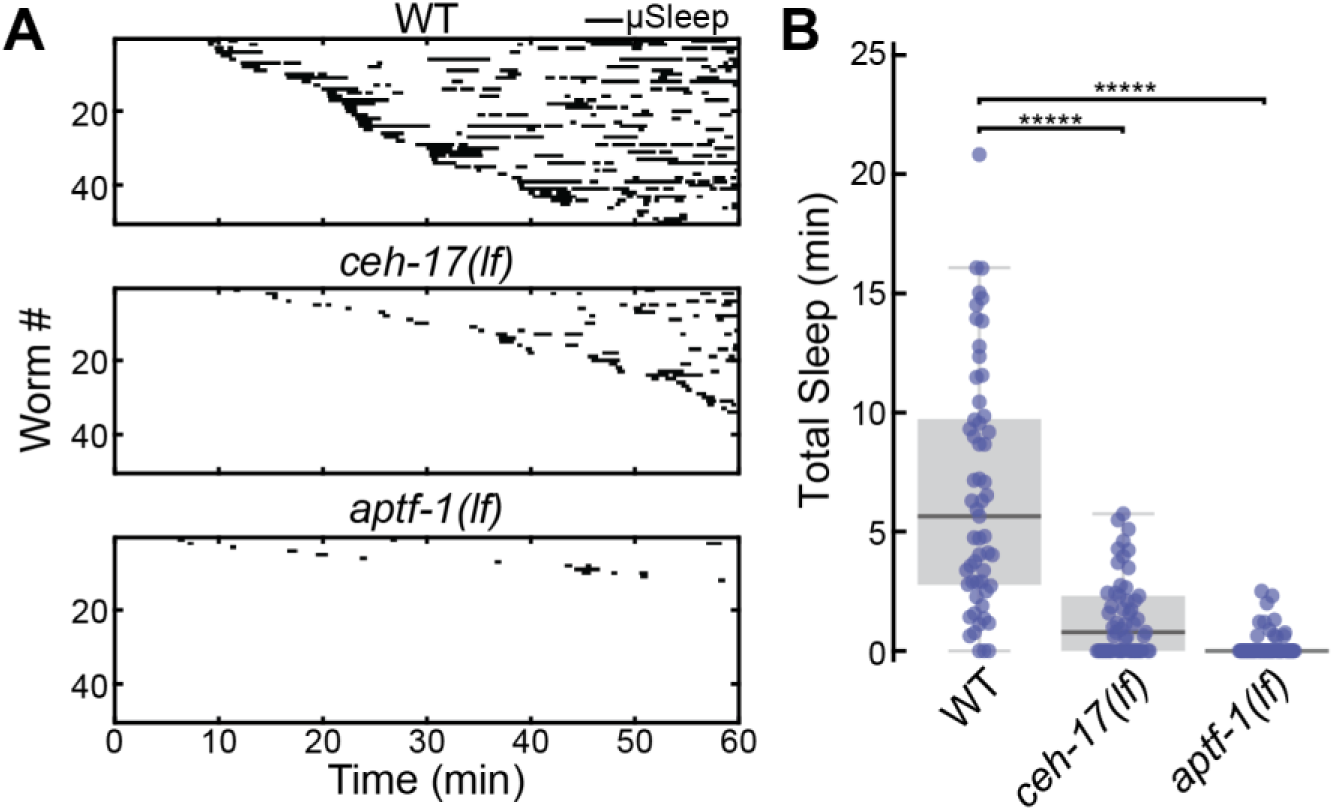
Microfluidic-induced sleep behavior depends on ALA and RIS neurons. *(A)* Raster plots of detected sleep bouts from WT, *ceh-17(lf)* and *aptf-1(lf)* animals. Only the top 50 animals showing the most total microfluidic-induced sleep are shown. *(B)* Both *ceh-17(lf)* and *aptf-1(lf)* show less total sleep than WT. The data suggest that microfluidic-induced sleep is strongly dependent on the ALA and RIS neurons. (WT n = 57, *ceh-17(lf)* n = 57, *aptf-1(lf)* n = 60; *****p < 0.00001 compared to WT, Kruskal-Wallis with a *post-hoc* Dunn-Sidak test).

**Figure 5.**
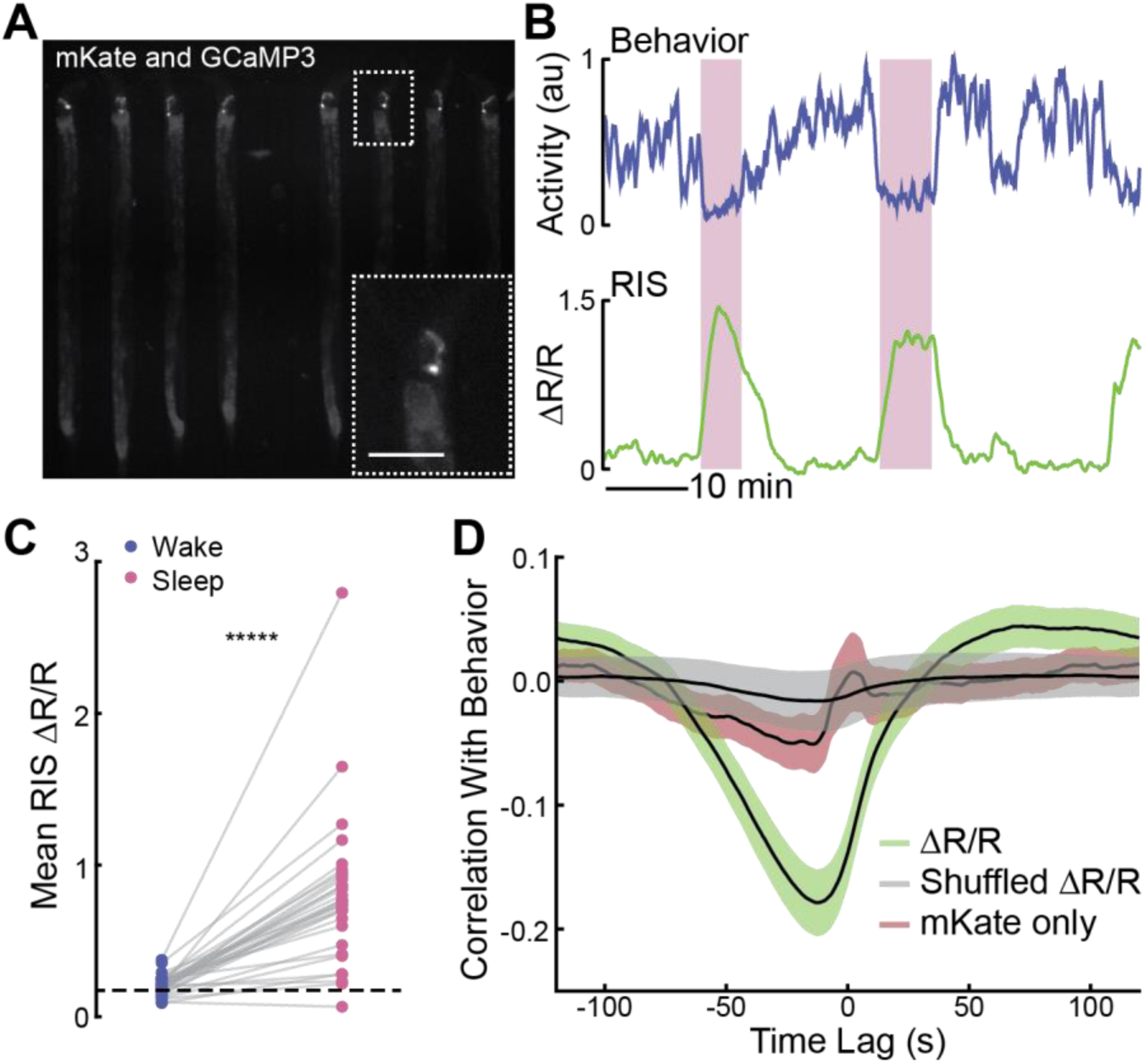
The RIS neuron is more active during microfluidic-induced sleep. *(A)* Fluorescent micrograph of the mKate channel, showing HBR1361 animals immobilized in 50 μm-wide chambers and expressing mKate and GCaMP3 in the RIS neurons. Inset is a zoom-in on a single RIS neuron (scale bar 50 μm). *(B)* Representative traces of animal behavioral activity (top) and RIS calcium activity (bottom). RIS activity dramatically increases during microfluidic-induced sleep bouts (shaded regions), opposed to the majority of worm brain activity (see Figure 4). *(C)* RIS is more active during microfluidic-induced sleep. We automatically detected sleep bouts across animals, calculated mean RIS activity during wake and microfluidic-induced sleep, and quantified mean RIS activity in each behavioral state. Dashed line shows the average RIS activity for animals that did not display a sleep bout. (Data points represent individual animals; n = 48 animals total, n = 31 animals exhibited at least one sleep bout, n = 17 animals did not sleep; *****p < 0.00001, paired t-test). *(D)* RIS ΔR/R activity negatively correlates with animal behavior. This correlation was not seen in shuffled ΔR/R data or when the mKate channel only was compared with behavior, indicating that RIS correlation with behavior is not a result of movement artifacts.

To further validate that microfluidic-induced sleep is a sleep state, we interrogated the role of two interneurons, RIS and ALA, known to dominate *C. elegans* developmental and stress-induced sleep, respectively, via at least partially distinct signaling pathways ^46, 64^. To test whether microfluidic-induced sleep is also dependent on these neurons, we compared the sleep phenotype of WT, *ceh-17(lf)* and *aptf-1(lf)* loss-of-function mutants (**Figure 3**). *ceh-17(lf)* shows defective axonal growth of the ALA neuron ^35^ and less stress-induced sleep ^40^. *aptf-1(lf)* lacks the transcription factor necessary for RIS to signal quiescence via the FLP-11 neuropeptide and shows defects in developmental sleep ^38, 39^. In our experiments, both *ceh-17(lf)* and *aptf-1(lf)* mutants showed more than 4.5 times less total microfluidic-induced sleep than WT animals (**Figure 3B**). Furthermore, while these behavioral data only represent sleep bouts lasting longer than 30 s to reduce false detections (see Methods), these results—and others throughout this report—were not dependent on this choice of analyses (**Supplementary Figure 3**). Together, these data validate that microfluidic-induced sleep is a *C. elegans* sleep state controlled by previously reported neural mechanisms.

Having established that microfluidic-induced sleep meets the criteria for *C. elegans* sleep and is hallmarked by a dramatic behavioral state transition, we also sought to confirm that these spontaneous behavioral transitions were accompanied by an underlying global-brain state transition ^33, 65^. Previous work in chemically-paralyzed animals used brain-wide calcium imaging to establish that the *C. elegans* nervous system, with the exception of a few neurons, transitions to a large-scale downregulation of neural activity during sleep ^33, 65^. Yet, this global brain activity has only been observed during *C. elegans* developmental sleep ^33^ and sleep induced by a 16 hr period of starvation ^65^, which are both significantly different conditions compared to microfluidic-induced sleep. For example, our microfluidic assays use adult animals and we typically observe quiescence within the first hour after removal from food (**Figure 1**, Methods).

Using whole-brain calcium-sensitive imaging we were able to establish that microfluidic-induced sleep is indeed associated with a global brain state transition (**Figure 4**). For these experiments we found that the microfluidic-induced sleep behavior has a distinct advantage compared to other whole-brain imaging experimental preparations. Typically, whole-brain calcium imaging in *C. elegans* exists in two paradigms: volumetric imaging of either freely-moving animals ^66–68^ or chemically-paralyzed animals confined in microfluidic chambers ^32, 33, 65, 69, 70^. While imaging freely-moving animals allows for simultaneous quantification of neural activity and behavior, tracking a moving brain during experiments ^66, 67^ and tracking the location of individual neurons during post-processing ^68^ remains challenging. Alternatively, chemically paralyzing animals simplifies experiments but does not provide a direct behavioral output during imaging ^32, 33, 65, 69, 70^. The major advantage of the microfluidic-induced sleep behavior for whole-brain imaging is that animals can be confined in microfluidic chambers, which facilitates imaging without the use of paralytics that could disrupt spontaneous sleep-wake transitions (see **Figure 1**, **Supplementary Movie 2**). To exploit this advantage, we developed an imaging protocol using a microfluidic chamber geometry similar to previous studies ^32, 33, 69, 70^ that partially-immobilizes animals but allows for enough movement to quantify animal behavior and detect microfluidic-induced sleep during whole-brain calcium imaging (**Figure 4**). By incorporating a period of animal habituation to blue excitation light, we were able to image continuously for 10 minutes using single-plane epifluorescence microscopy and capture spontaneous sleep-wake transitions in approximately 25% of animals (see Methods). Imaging a single 2D-plane allowed for quantifying both average ganglia activity and the activity of several individual neurons (**Figure 4B**, **Supplementary Movie 6**).

**Figure 4.**
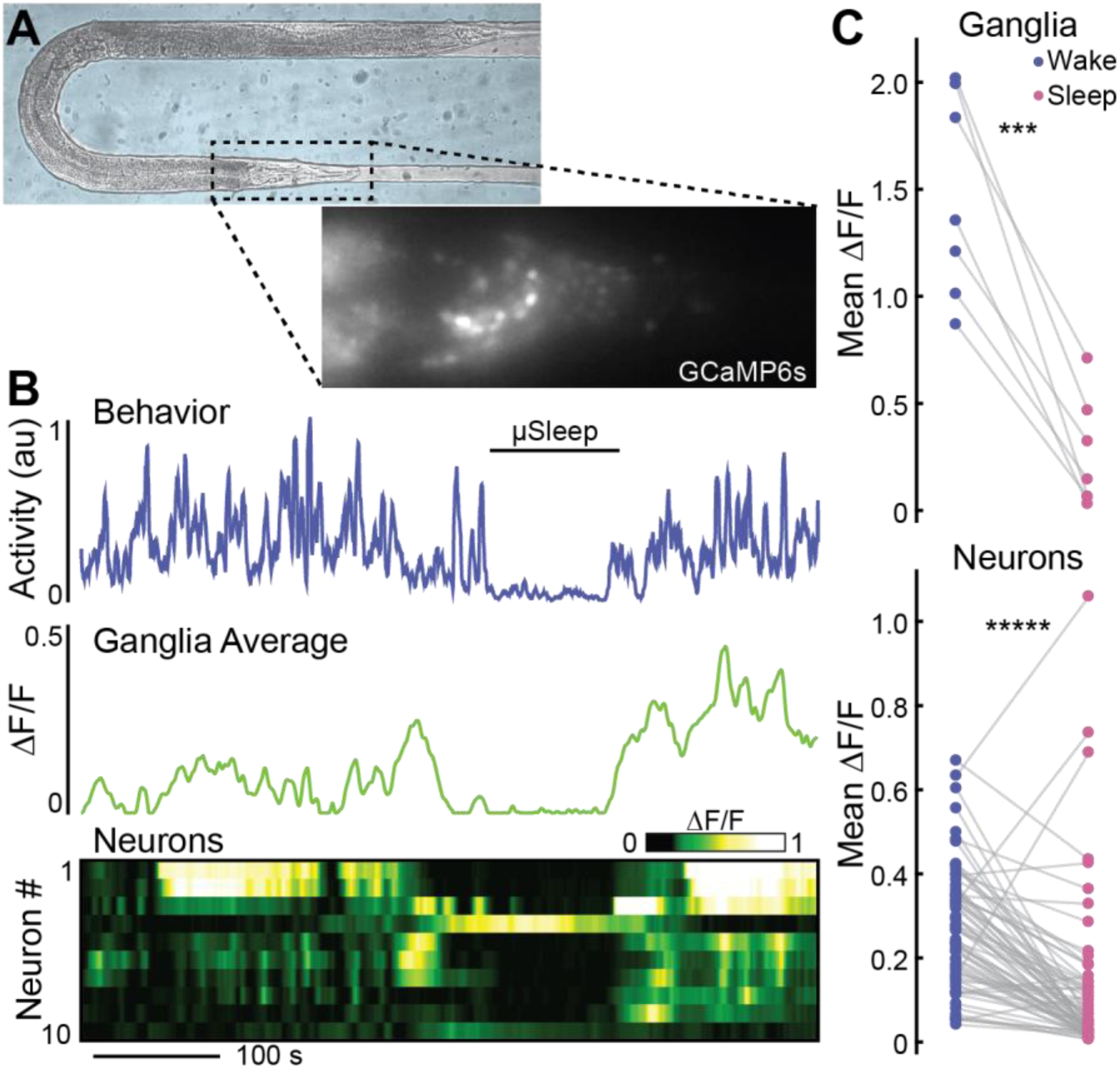
A global-brain state transition governs *C. elegans* microfluidic-induced sleep. *(A)* Adult animal immobilized in a microfluidic chamber tailored for whole-brain imaging. *(Inset)* Single-plane epifluorescence image of an animal with pan-neuronal expression of GCaMP6s. *(B)* Representative animal shows behavioral quiescence correlates with less neural activity. *(Top)* Behavioral activity trace quantified by tracking the motion of ten individual neurons (see Methods). *(Middle)* Average fluorescence across the whole worm head ganglia. *(Bottom)* Activity of ten individual neurons show a clear brain-state transition and less neural activity during microfluidic-induced sleep. *(C)* Using only behavioral activity, we identified quiescent bouts then quantified neural activity during sleep and wake. During microfluidic-induced sleep, animals exhibited a large-scale downregulation of neural activity across both the entire ganglia and most individual neurons. The neurons chosen for analysis were randomly selected from the field-of-view and are not necessarily the same neurons across every animal. (n = 7 animals; 10 neurons were tracked per animal; ***p < 0.001, *****p < 0.00001, paired t-test).

With this imaging protocol, we observed a distinct correlation between animal behavior and neural activity (**Figure 4B**, **Supplementary Movie 6**). During microfluidic-induced sleep both the average ganglia activity and the activity of individual neurons dropped significantly (78 ± 10% and 55 ± 10%, respectively) (**Figure 4C**). However, as expected from previous work^33, 38, 39, 43, 65^, we observed that some neurons actually increased in activity during microfluidic-induced sleep (**Figure 4C**). Single-neuron imaging confirmed that the RIS neuron, which has been proposed as a sleep-promoting neuron for multiple *C. elegans* sleep-like states ^38, 39, 43^, indeed showed more than a two-fold increase in calcium activity during microfluidic-induced sleep and strongly correlated with low-behavioral activity (**Figure 5**). These results further support the claim that microfluidic-induced sleep is a *C. elegans* sleep behavior controlled by sleep-promoting circuits and corroborate previous reports that a unique brain state governs *C. elegans* sleep ^33, 65^. Furthermore, these results show that microfluidic-induced sleep behavior is an advantageous behavioral paradigm that facilitates whole-brain imaging while measuring animal activity without the need for chemically induced paralysis.

While the combination of behavioral and calcium-imaging data shows that microfluidic-induced sleep is hallmarked by a spontaneous brain *and* behavioral state transition (**Figure 4**) regulated by sleep-promoting neurons like RIS and ALA (**Figure 3**, **Figure 5**), we also wanted to understand what neural circuits upstream of these neurons are involved in regulating microfluidic-induced sleep. Based on our observation of distinct phenotypes when comparing swimming animals in WorMotel to swimming animals in microfluidic chambers (**Figure 1**), we hypothesized that *C. elegans* sensory circuits detect features unique to the microfluidic environment and in turn drive sleep behavior. To elucidate the cues regulating microfluidic-induced sleep, we used the versatility of microfluidics to perform assays under a variety of conditions and compared the microfluidic-induced sleep phenotype to the baseline behavior observed in a large microfluidic device (**Figure 6A**). We defined our baseline experimental conditions as: 20 °C cultivation temperature (*Tc*); animals are transferred directly from seeded nematode growth media (NGM) into the microfluidic device; the microfluidic media (M9 buffer) contains no food source; the temperature during imaging is 22 °C.

**Figure 6.**
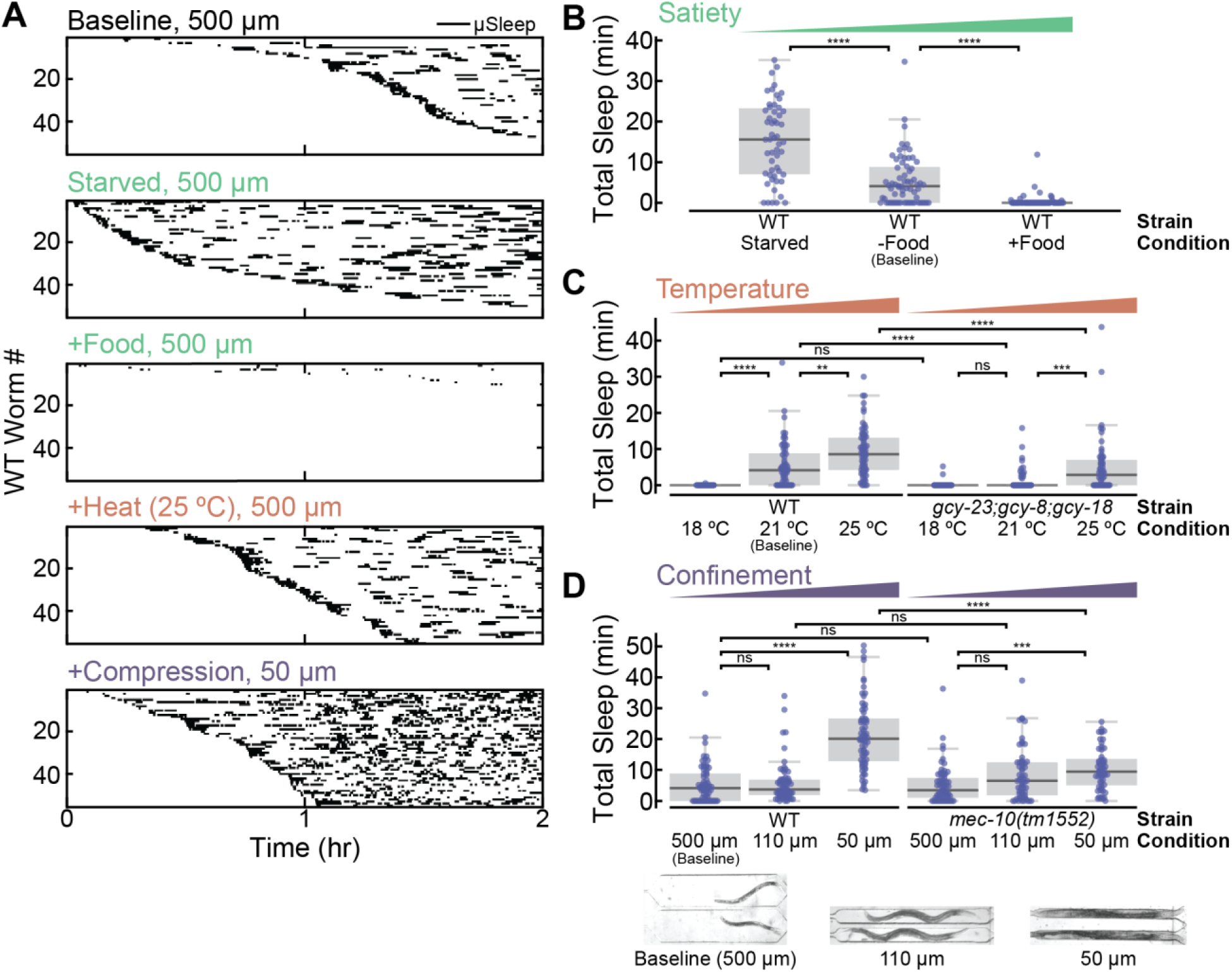
microfluidic-induced sleep is regulated by satiety and multiple sensory circuits. *(A)* Detected sleep bouts for WT animals in several experimental conditions. Raster plots show detected sleep bouts during a 2 hr imaging period. “Baseline” indicates the standard experimental conditions: 500 μm chamber width, no food in the buffer, and a 22 °C temperature. “Starved” indicates animals that were starved prior to the assay. “+Food” indicates conditions in which *E. coli* OP50 was added into the buffer during recordings. “+Heat” indicates imaging conditions where the temperature was raised to 25 °C. “+Compression” indicates that animals are partially immobilized in 50 μm-wide chambers. See micrographs under (D) for chamber geometries. In all cases, the sleep phenotype varies dramatically from the baseline. Only the 55 animals that displayed the most sleep are plotted for clarity. *(B)* Total WT sleep under varying satiety conditions. As satiety increases from “Starved” to “+Food,” animals exhibit less microfluidic-induced sleep (from left to right on the plot the number of animals n = 55, 68, 67). *(C)* Total microfluidic-induced sleep under varying temperature conditions. Increasing temperature increases total microfluidic-induced sleep for WT animals. Thermosensory-defective mutants show the same microfluidic-induced sleep phenotype as WT at 18 °C, but significantly less sleep as 22 °C and 25 °C, indicating that thermosensory input can act to drive or suppress microfluidic-induced sleep (from left to right on the plot the number of animals n = 37, 68, 71, 41, 67, 60) *(D)* Total sleep under different confinement conditions. Micrographs show chamber geometries. When WT animals are confined in smaller chambers, they only show an increase in total microfluidic-induced sleep when slightly compressed in 50 μm-wide chambers. *mec-10(tm1552)* mutants show an identical phenotype to WT in 500 μm and 110 μm chambers, but dramatically reduced sleep compared to WT when compressed. These results suggest that nociceptive input to mechanosensory neurons regulates microfluidic-induced sleep (from left to right on the plot the number of animals n = 68, 64, 69, 66, 64, 57). (ns = not significant, ** p < 0.01, ***p<0.001, ****p < 0.0001; Kruskal-Wallis with a *post-hoc* Dunn-Sidak test).

We first tested how food availability affects the microfluidic-induced sleep phenotype. The presence of food is known to change the swimming-induced quiescence phenotype ^52, 53^, high quality food induces quiescence ^71^, and extreme starvation also leads to sleep-wake switching ^43, 65^. Therefore, we hypothesized that the internal satiety state is a significant regulator of microfluidic-induced sleep. To test this hypothesis, we assayed three satiety states in WT animals: “Starved” animals were starved for 2 hr before being loaded into a chambers with no food; “Baseline” animals were loaded directly from a NGM food source to a microfluidic device with no food; and “+Food” animals were loaded from a NGM food source into a microfluidic device with food dissolved in the buffer. As expected, we observed less total microfluidic-induced sleep per worm as satiety increased from “Starved” (15 ± 2 min, mean ± sem) to “Baseline” (5.4 ± 0.9 min) to “+Food” (0.4 ± 0.2 min) (**Figure 6A-B**). We also observed that only in the “Starved” condition did animals show measurably lower behavioral activity during wakefulness (**Supplementary Figure 4A**). These results show that satiety is a strong regulator for microfluidic-induced sleep, which may have evolved to optimize the tradeoff between energy conservation and the search for food.

Other well-known drivers of sleep are noxious environmental stressors, which we also tested in the context of our microfluidic chambers. Heat shock at temperatures greater than 30 °C is a common method to induce cellular stress and *C. elegans* sleep ^34, 40, 44, 45, 54, 72^. Although our standard microfluidic-induced sleep assays were conducted at only 22 °C, we investigated whether these small changes from *Tc* could be a possible environmental regulator of quiescence. Indeed, we discovered that increasing the assay temperature to 25 °C, which is still well below commonly used temperatures for heat-shock (30-40 °C), increased the total observed sleep 3.5- fold (**Figure 6A, C**). Cooling the device to 18 °C (i.e. below *Tc*) had the opposite effect and nearly abolished all microfluidic-induced sleep bouts (**Figure 6C**). To show that these changes in phenotype are mediated by *C. elegans* thermosensation, we tested triple-knock-out *gcy-23(nj37);gcy-8(oy44);gcy-18(nj38)* mutants, which show disruption to thermosensation that is specific to the AFD neurons, the primary temperature-sensing neurons in *C. elegans* ^73, 74^. In AFD-defective animals we observed no significant change in total microfluidic-induced sleep when raising the temperature from 18 °C to 22 °C (baseline). Under the same conditions, WT animals showed a 540-fold increase in total microfluidic-induced sleep (**Figure 6C**).

Furthermore, thermosensory-defective mutants displayed 80% less microfluidic-induced sleep at 22 °C (baseline) and 45% less microfluidic-induced sleep at 25 °C when compared to WT animals (**Figure 6C**). These results suggest that AFD neurons transduce changes in environmental temperature and drive downstream sleep-promoting circuits. In addition, temperature had no effect on wake-state behavioral activity for either WT or mutant animals (**Supplementary Figure 4B**). These results show that temperature changes significantly less than those typically used for heat shock can dramatically change sleep dynamics. In addition, this is the first indication that thermosensory input can act bidirectionally to either promote or suppress *C. elegans* quiescence.

The final environmental factor we investigated was animal confinement. Under baseline conditions, animals swim in large 500 μm-width chambers (**Figure 6D**, inset). We found that confining animals to a smaller 110 μm-width chamber, where the primary motion is crawling-like (**Figure 6D** inset) leads to the same amount of total microfluidic-induced sleep as the baseline chambers (**Figure 6D**). However, when we partially compressed WT animals in small 50 μm-width chambers, we observed nearly a 4-fold increase in total sleep compared to baseline (**Figure 6A, D**). We hypothesized that this change in phenotype due to immobilization was at least partially mediated by mechanosensory circuits. For example previous reports showed that touch-defective mutants *mec-10(tm1552)* have a decreased ability to sense spatial patterns in microfluidic arenas ^75^. We tested *mec-10(tm1552)* as well and found that when minimal mechanical stress was present (i.e. 500 μm and 110 μm width chambers), total microfluidic-induced sleep remained insignificantly unchanged compared to WT (**Figure 6D**). However, when *mec-10(tm1552)* animals were compressed, total animal microfluidic-induced sleep decreased by a factor of two compared to WT animals under the same conditions (**Figure 6D**), indicating that mechanosensory pathways are necessary for restraint-induced microfluidic-induced sleep. While animal restraint indeed changed the measured behavioral activity, these differences in sleep between WT and *mec-10* animals cannot be explained by changes in behavioral activity during wakefulness (**Supplementary Figure 4C**). In addition, we also tested *mec-4(e1611)* mutants, which are defective in response to gentle touch, but remain sensitive to harsh touch ^76^. These animals showed no significant difference in total microfluidic-induced sleep compared to WT when immobilized (**Supplementary Figure 5**), indicating that nociceptive mechanosensory pathways are required to drive microfluidic-induced sleep. Together, these results show for the first time that *C. elegans* microfluidic immobilization induces sleep behavior through harsh mechanosensory pathways.

Given the dependence of microfluidic-induced sleep on satiety, temperature, and mechanical stress (**Figure 6**), we hypothesized that each of these factors could act as stressors that contribute to a slow-onset form of stress-induced sleep. Previously, studies of stress-induced sleep in *C. elegans* have reported acute and rapid sleep responses to noxious external stimuli ^34, 40, 44, 45, 54, 72^, leading us to investigate if the slow-onset microfluidic-induced sleep state could be weaker form of stress-induced sleep. To test this hypothesis, we assayed DAF-16::GFP animals in the identical environmental conditions shown in **Figure 6**. The Fork head transcription factor DAF-16 regulates longevity, metabolism, and some stress responses in *C. elegans* ^77–79^. While normally diffuse in the cytoplasm, under unfavorable environmental conditions (e.g. heat shock), DAF-16::GFP localizes to the nucleus ^79^. Thus DAF-16::GFP puncta formation can serve as a proxy for *C. elegans* stress under some conditions. We quantified puncta formation in different environmental conditions and observed that DAF-16::GFP localization correlated with microfluidic-induced sleep phenotypes in only some cases (**Supplementary Figure 6**). For example, compared to baseline animals, animals assayed at 25 °C showed greater than a 250% increase in the number of cumulative puncta and animals assayed with food showed almost zero puncta formation (**Supplementary Figure 6**), which is in agreement with previous reports of stress responses ^79, 80^. These results support the hypothesis that 25 °C and the lack of food in the microfluidic devices increases animal stress and upregulates microfluidic-induced sleep. However, animals partially immobilized in 50 μm-wide chambers showed no significant difference in the number of DAF-16 puncta compared to Baseline (**Supplementary Figure 6**), despite displaying dramatically more total microfluidic-induced sleep (**Figure 6**). These results suggest that animal stress may not completely explain the changes in microfluidic-induced sleep we observe during environmental manipulations.

In addition to environmental elements like mechanical compression and temperature that change sleep, we ruled out the possibility that O2 depletion, CO2 buildup, or the buildup of other unknown substances dramatically affect microfluidic-induced sleep (**Supplementary Figure 7A-C**). To rule out these factors performed two different experiments. In one, we replaced the static buffers (used for all other experiments) with a gentle buffer flow that is expected to stabilize gas concentration levels and remove any animal byproducts. Surprisingly, we observed that the gentle flow led to more sleep (**Supplementary Figure 7A-C**). In another experiment, we considered the possibility that the small volume of liquid in different microfluidic chambers could play a role in driving more sleep through the buildup of gasses or other products. To test this hypothesis, we developed a geometry that maintains the same animal restraint but varies the fluidic volume in the worm chambers (**Supplementary Figure 7D**). The total sleep in these chambers was not significantly different (**Supplementary Figure 7E-F**). These experiments suggest that indeed the mechanical environment is a much stronger driver of sleep than fluidic volume, and not the buildup of metabolic products.

Finally, we also ruled out the possibility that the process of loading animals into the microfluidic device or changing the fluidic environment leads to a stress response that drives stress-induced sleep. We fabricated large microfluidic chambers that are expected to mimic an open volume of liquid (1200 μm long, 500 μm wide, 90 μm tall), and observed that animals rarely exhibited sleep behavior despite undergoing the same loading process (**Supplementary Figure 7G-I**). Therefore, the stress associated with the loading process and change in the fluidic environment alone is not sufficient to drive stress-induced sleep. In fact, reducing this chamber height to 55 μm reestablished the sleep behavior, further strengthening our claims that the mechanical environment is a strong driver of microfluidic-induced sleep (**Supplementary Figure 7G-I**).

Future work will continue to interrogate how sensory mechanisms regulate sleep and arousal. Overall, our environmental manipulations conclusively show that the animal satiety state, the mechanical environment, and the local temperature strongly regulate the microfluidic-induced sleep phenotype. Thus, microfluidic-induced sleep is a multisensory-regulated sleep behavior that is also strongly modulated by internal physiological factors.

We envision microfluidic-induced sleep as a platform to understand the neural circuits and correlates driving brain-state transitions. To highlight this potential, we performed additional experiments to record from more individual neurons during whole-brain imaging. As shown in previous works, tracking individual neurons in a moving *C. elegans* brain is a laborious and challenging process ^66, 68^. Following examples from other groups ^32, 33, 65, 69, 70^, we circumvented this challenge by chemically paralyzing animals, performing z-scanning, and capturing the activity of 40-100 neurons simultaneously during volumetric imaging (**Supplementary Figure 8**). In a fraction of animals, we observed brain-states that resembled what would be expected from a sleeping animal (decrease in calcium activity in all but a few neurons, see **Figure 4**).

However, in this paralyzed preparation we cannot unequivocally claim that these states correspond to sleep because we cannot correlate the calcium imaging with animal behavior. Nevertheless, our data is consistent with our expectations for how paralysis should affect the sleep state given the role of mechanoreception. We found that the probability of observing a putative sleep state during a 10 minute experiment dropped from a rate of ∼25% in behaving animals to <5% in paralyzed animals. This result is consistent with our conclusion that mechanosensation of the environment facilitated by worm movement is a primary driver of microfluidic-induced sleep. Though we are limited here by a relatively slow imaging rate (< 3 vol/s) and poor z-sectioning with epifluorescence imaging, these experiments show a path towards using microfluidic-induced sleep to understand how neural circuits drive spontaneous brain-state transitions. Future directions should explore high-resolution rapid volumetric imaging techniques ^81^ to measure both the calcium activity and neuron location (e.g. using GCaMP6s and RFP ^66–68^), to not only capture brain-wide activity but also enable single-neuron mapping to the *C. elegans* connectome.

## Discussion

Here, we described a spontaneous *C. elegans* brain and behavioral state transition that is unique to microfluidic chambers, which facilitates whole-brain imaging and precise regulation of the environment. microfluidic-induced sleep regulation appears to be at least partially linked to a slow-onset form of stress-induced sleep (**Supplementary Figure 6**), and may also be related to swimming-induced quiescence ^49, 51, 52^. Despite these similarities, the behavioral dynamics of quiescence that we measured is quantitatively distinct from these previously reported worm quiescent behaviors (**Figure 1**), although it is possible that changes in sleep onset time and bout duration in microfluidics is affected by collisions with the chamber walls or other physical cues. We showed that during microfluidic-induced animals exhibit many behavioral properties of sleep, including a stereotyped posture (**Figure 2A**), reversibility (**Figure 2B**), a decreased response to sensory stimuli (**Figure 2C-D**), and homeostatic rebound (**Supplementary Figure 2**). In addition, this quiescence behavior is dependent on known *C. elegans* sleep-promoting neurons, ALA and RIS (**Figure 3**). Thus, microfluidic-induced sleep meets the precedent for sleep behavior^24, 40, 46^. In addition, we found that thermosensory input via the AFD neuron acts as a bidirectional controller of sleep; cooler temperatures promote wake, while warmers temperatures promote sleep (**Figure 6**). Furthermore, we showed for the first time that animal restraint can act through mechanosensory pathways to drive sleep behavior (**Figure 6**). Finally, a dramatic brain-state transitions and global downregulation of neural activity, with the exception of a few neurons, underlies microfluidic-induced sleep behavioral transitions (**Figure 4**, **Figure 5**, **Supplementary Figure 8**).

Importantly, our results show the drastic effects of microfluidic environments on *C. elegans* behavior. Microfluidic assays are commonly used to study nematode biology including, aging, behavior, and neurobiology ^82–84^. Sleep in *C. elegans* is also commonly studied by confining animals to microfluidic chambers ^52, 53, 56, 85–87^. Our data show that the conditions of these micro-environments can strongly affect the sleep phenotype and thus the microfluidic chambers should be carefully designed and their effects should be considered when interpreting the results of these experiments. However, although we observe changes in sleep behavior in the context of microfluidics—and most notably during increased confinement—quiescence behavior may increase in standard agar environments as well due to changes in animal confinement. For example, we hypothesize that experiments with agarose-based confinement chambers ^86, 88^ would observe the similar changes in behavior to those reported here.

Furthermore, it is not difficult to imagine *C. elegans* has evolved to sense small changes in temperature, food, and confinement and use these environmental cues to drive quiescence behavior and a downregulation of neural activity. One unifying hypothesis for *C. elegans* microfluidic-induced sleep regulation could be rooted in energy conservation. Factors that decreased microfluidic-induced sleep, such as food and cooler temperatures, may indicate favorable conditions for active roaming behavior. Conversely, factors that increased microfluidic-induced sleep, such as elevated temperatures and animal restraint, may indicate more harmful environments in which roaming and expending energy is not an optimal survival strategy. Furthermore, the changes we made to the environment may change the animal’s metabolic state. *C. elegans* metabolism is known to increase with increasing temperature ^89^. Additionally, one can imagine that microfluidic immobilization (**Figure 6**) drives an escape response, leading to increased muscle activation and high energy expenditure. Therefore, it is possible that all changes in microfluidic-induced sleep phenotypes we observed were rooted in optimizing energy use by balancing the between tradeoff between expending energy to move and the expected value of finding food. Future work is needed to continue to elucidate these mechanisms using genetic mutants, different culture conditions, and combining sensory stimuli with food availability. Overall, the ability to control food availability, temperature, and mechanosensory feedback provides a valuable platform to study how quiescent behavior may serve as an evolutionary advantage and successful survival strategy.

Many questions remain with respect to *C. elegans* sleep and the role of sensory inputs in driving sleep behavior. For example, sleep-promoting neurons such as ALA and RIS drive behavioral quiescence, but what ensemble activity occurs upstream of these circuits? The interaction between multiple sensory inputs, the ALA/RIS neurons, and diverse behavioral outputs are likely highly complex, non-linear systems involving multiple pathways and neuromodulators ^34, 38, 43–45, 53^. Future work will help answer these questions and determine the relationship between ALA and RIS and whether these neurons are critical for driving sleep in all the environmental conditions tested in this report (**Figure 6**). Furthermore, ALA and RIS themselves are known to respond to mechanical stimulation ^86, 90^, potentially providing a mechanism for why animal confinement drives sleep behavior. These “sleep-driving” neurons that also act as sensory neurons are a prime example of the multimodal nature of the C. elegans nervous system, which adds another element of complexity when dissecting the sensory and sleep circuits involved in environmentally-driven sleep.

In a broader context, we have identified microfluidic-induced sleep as behavior in which external inputs can be tightly regulated to manipulate animal behavior. Therefore, this sleep behavior provides a testbed for interrogating the biological principles that regulate brain and behavioral state transitions and investigating how these principles adapt in changing environments and physiological states. These studies are made possible by the combination of whole-brain imaging and behavioral recordings with simultaneous control of the microfluidic environment. Future opportunities include volumetric whole-brain imaging ^66–68^ that could capture the activity of nearly every neuron in the worm brain to elucidate how ensemble activity drives state transitions. Combining these imaging methods with high-throughput microfluidic assays and dynamic environmental control will lead to a large library of simultaneous behavioral and brain-wide data under many experimental conditions. These data, combined with computational techniques, would provide a rich resource to understand how many circuits across the *C. elegans* nervous system regulate brain- and behavioral states.

## Acknowledgments

We thank the Fang-Yen lab for providing the WorMotel device. Henrik Bringmann provided the HRB1361 strain for RIS imaging and HBR1374 strain for optogenetics. We thank Anne Hart, David Raizen and Henrik Bringmann for helpful discussions. Several strains were provided by the CGC, which is funded by NIH Office of Research Infrastructure Programs (P40 OD010440). D.L.G. is funded by the National Science Foundation (NSF) Graduate Research Fellowship Program 1842494 and a training fellowship from the Keck Center of the Gulf Coast Consortia on the NSF IGERT: Neuroengineering from Cells to Systems 1250104. D.L.G. and J.Z. are funded by the Smalley-Curl Institute’s Student Training for Advising Research program. We also thank the Rice Shared Equipment Authority where devices were fabricated.

## Author contributions

D.L.G. conceived, performed, and analyzed experiments. J.Z. performed whole-brain imaging experiments. B.F. performed experiments. J.T.R. directed the research. D.L.G. and J.T.R. co-wrote the paper. All authors read and commented on the manuscript.

## Competing financial interests

The authors declare no competing financial interests.

## METHODS

*C. elegans* Strains and Maintenance

All *C. elegans* strains were maintained at 20 °C on standard Nematode Growth Medium (NGM) seeded with *E. coli* OP50 as the food source. All experiments were performed with day-1 adult animals at 21-22 °C unless otherwise stated. The strains used for each experiment are as follows

**Figure 1:** N2

**Figure 2:** N2

**Figure 3:** N2, IB16 *ceh-17(np1)* I; HBR227 *aptf-1(gk794)* II

**Figure 4:** AML32 (wtfIs5 [prab-3::NLS::GCaMP6s; prab-3::NLS::tagRFP])

**Figure 5:** HRB1361 (goeIs304[pflp-11::SL1-GCaMP3.35-SL2::mKate2-unc-54-3’UTR, unc-119(+)])

**Figure 6:** N2, IK597 gcy-23(nj37);gcy-8(oy44);gcy-18(nj38) IV, ZB2551 mec-10(tm1552)

**Supplementary Figure 1.** N2

**Supplementary Figure 2:** N2, HBR1374 goeIs307[pflp-11::ArchT::SL2mKate2-unc-54-30UTR,unc-119(+)]; goeIs304[pflp-11::SL1-GCaMP3.35-SL2::mKate2-unc-54-30UTR, unc-*119(+)]*

**Supplementary Figure 3:** N2, IB16 *ceh-17(np1)* I; HBR227 *aptf-1(gk794)* II

**Supplementary Figure 5:** N2, CB1611 *mec-4(e1611)*

**Supplementary Figure 6:** TJ356 (zIs356 [daf-16p::daf-16a/b::GFP + rol-6(su1006)])

**Supplementary Figure 7:** N2

**Supplementary Figure 8:** AML32 (wtfIs5 [prab-3::NLS::GCaMP6s; prab-3::NLS::tagRFP])

### Microfluidic Device Fabrication

Standard photo- and soft-lithography techniques were used to fabricate microfluidic devices ^84^. Microfluidic geometries were custom designed in CAD software. Most photomasks were transparencies (CAD/Art Services Inc.), but glass photomasks (Front Range Photomask) were used for higher-resolution devices (**Supplementary Figure 7**). All master molds were fabricated using SU-8 2075 (MicroChem). The SU-8 height for worm behavioral channels was 75 μm (spin: 20 s – 500 rpm, 30 s – 3000 rpm), but we used a height of 50 μm (spin: 20 s – 500 rpm, 30 s – 4000 rpm) for whole-brain imaging devices to further constrict animal movement. We used polydimethylsiloxane (PDMS) Sylgard for all microfluidic chips. All behavioral chips were double-layer devices to incorporate push-down valves for sealing the chamber entrances or delivering mechanical stimuli. The bottom worm layer (20:1 ratio, spin: 930 rpm for 30s) was bonded to the upper valve layer (10:1 ratio) using a 30 s exposure to oxygen plasma (200 W, 330 mTorr), then baked together for at least 12 hr. The PDMS devices were permanently bonded to either a standard glass slide for behavior, or a 300 μm-thick quartz wafer (NOVA Electronics Materials) for whole-brain imaging.

### Behavioral Quantification and Sleep Detection

To quantify *C. elegans* activity, we used the previously described method of frame-by-frame subtraction ^85^ using custom MATLAB (MathWorks) scripts. Not only is frame-by-frame subtraction less computationally expensive than other metrics such as posture or nose-tracking, but we found overall that this technique is less prone to the noise associated with small errors in tracking specific *C. elegans* body points. This method subtracts consecutive frames from one another, then counts the number of pixels that substantially change value. Worm movement leads to a large number of pixels that change from frame-to-frame. Quiescent animals, which move very little, lead to few pixels that change values from frame-to-frame. We performed frame-by-frame subtraction, drew regions-of-interest around each animal, then counted the number of pixels in the ROI that changed by a value greater than 30 (a number well above the noise level of the CMOS sensor). This yields a raw activity trace for each animal. We also used standardized methods to normalize activity traces ^52, 53^, which accounts for changes in brightness across the field-of-view (FOV), and allowed us to set a consistent sleep detection threshold across populations. In most cases, we smoothed activity traces across 20 s, then normalized to the top 95^th^ percentile, yielding a normalized activity trace for each animal with values approximately between 0 and 1. For analysis that required finer timescales on the order of less than 20 s (Figure 2A-C), we did not smooth activity traces (see “Reversibility and Decreased Response to Stimuli” subsection).

Once we normalized the activity trace, we thresholded the data to detect sleep. The threshold depended on the microfluidic geometry. For example, in large microfluidic chambers (Supplementary Movie 1), movement could still be detected during microfluidic-induced sleep as animals drifted across the chamber. In smaller microfluidic geometries (Supplementary Movie 2), worm activity was already significantly constrained, so a stricter threshold was needed to detect sleep. We determined thresholds by manually scoring 20 sleep bouts, calculating the mean activity during those bouts, then doubling the mean activity. The thresholds used for each geometry were: WorMotel: 0.15, 500 μm microfluidic chambers: 0.15, 110 μm microfluidic chambers: 0.08, 50 μm chambers: 0.06. A minimum sleep bout time of 30 s was used to reduce false detections. In addition, during a sleep state brief animal twitching lasting less than 15 s was not counted as a wake period.

### Standard Microfluidic Behavioral Assays

Unless otherwise stated, behavioral assays took place on an enclosed AmScope SM-2T-LED stereo microscope. Red transparency was used to filter out LED wavelengths that potentially affect animal behavior. Imaging was performed at 3 fps with either a Point Grey Grasshopper (GS3-U3-23S6M-C) or Basler Ace (acA1920-40um) CMOS cameras, which have nearly identical sensor specifications. While the experiments were typically carried out in a room held at 20 °C, the heat from the LED raised the microfluidic device temperature to 21-22 °C (the “Baseline” temperature in Figure 6).

M9 buffer was used for all experiments, with no food added unless otherwise stated. During standard experimental conditions, we used a hair pick to transfer day-1 adults directly from seeded NGM into the buffer of an open syringe cap. We then suctioned animals into Tygon tubing that led to the microfluidic chambers. The process of loading 6-12 animals into the chambers (depending on the geometry used) typically took less than 5 min. A push-down valve closed off the entrance of all worm chambers to prevent animals from escaping during imaging. After use each day, devices were flushed with ∼6 mL of DI water, sonicated for 10 min, flushed again with DI water, boiled for 15 min, and flushed a final time before storage at ∼80 °C overnight.

### WorMotel Assays

A 48-well WorMotel molded from PDMS was provided by the Fang-Yen Lab ^56^. Only 12 wells were used simultaneously. Prior to use, the PDMS was exposed to oxygen plasma for 30 s to make the surface hydrophilic. Each well was then filled with 8 μL of M9 buffer. Using a hair pick, we removed individual animals from seeded NGM, washed them in an M9 droplet, then transferred them into the WorMotel wells. For imaging, the PDMS was inverted and reversibly sealed on a glass slide during imaging (see Churgin *et al.* 2017). The glass slide was treated with Rain-X to prevent condensation. The device was illuminated obliquely with three AmScope Goose-neck LEDs that were filtered with red transparency. As with microfluidic devices, the temperature during imaging was 21-22 C. We imaged from below at 3 fps with a Point Grey Grasshopper (GS3-U3-23S6M-C).

### Heat-Shock

Device setup and animal loading was performed as previously stated (see Standard Microfluidic Behavioral Assays subsection). However, the device was preheated to 30 °C with two Peltier heaters placed on each side of the glass slide. Heat was applied for the initial 30 min of imaging, after which the current through the Peltier heaters was reversed such that the device was rapidly brought to the standard 22 °C. This heat-shock protocol was adapted from that used for agar plates in Hill *et al.* 2014.

### Posture/Body Curvature Analysis

Similar to previous reports^57, 91^, Custom MATLAB scripts were written to quantify body curvature in the “Large μFluidic” data set from Figure 1 (where animals are confined to microfluidics, but have enough room to swim). We limited our analyses to the first two hours of imaging, when a minimal number of eggs are present. Many eggs in the chamber can make it computationally difficult to distinguish between eggs and the animal body. For every video frame, we subtracted the background (an image of the chambers with no animals present) and increased the frame contrast (MATLAB imadjust). These preprocessing steps yielded a high contrast between the animal bodies and image background. We then binarized the video (MATLAB im2bw), removed small objects (MATLAB bwareaopen), smoothed the binarized frame (MATLAB imgaussfilt), and skeletonized the smoothed frame (MATLAB bwmorph), and removed spurious pixels (MATLAB bwmorph). These steps gave skeletonized versions of the worm bodies. We used these skeletons to calculate the average body length and placed 20 equally spaced points along the centerline. Occasionally (<5% of frames), this centerline fit performed poorly (e.g. animals completely curl on themselves), but this could be detected by calculating the body length in every frame. Frames where the body length was less than 80% of the average were not used to calculate curvature. From the centerline points, we fit a circle to every 3 adjacent points, yielding 18 curvature values along the body (the inverse of the circle radii). These 18 curvature values were averaged and normalized to the body length to give a dimensionless value for the mean animal body curvature in every frame. For every animal, we then smoothed the curvature data over 20 seconds and found the average curvature during sleep and wakefulness.

### Reversibility and Decreased Response to Stimuli

Reversibility assays were conducted on an inverted Nikon microscope and performed with 50 μm-wide microfluidic chambers (**Supplementary Movie 3**). Animals were loaded into the chambers and left for 30 min. Between 30 and 60 min, upon visual confirmation of sleep, we initiated a protocol of 60 s of imaging at 10 fps. At 30 s, a strong light stimulus (460 nm light at 5 mW/mm^2^) was presented for 5 s to awaken animals. Only animals that were asleep for the full 30 s prior to the light stimulus were kept for analysis.

To test for a decreased response to stimuli, we fabricated 110 x 1100 um microfluidic chambers to confine animals (**Supplementary Movie 4-Supplementary Movie 5**). In addition to the push-down valve used to keep animals in each chamber, we incorporated two push-down valves used for mechanical stimulation. The two valves ensured that animals were almost always in contact with at least one valve during pressurization. For strong and weak stimulation, we used a valve pressure of 30 and 15 psi, respectively. 7 groups of animals were used for weak stimulation and 4 for strong stimulation; each group consisted of 8-12 animals. During the hour-long assay, animals received a 0.5 s valve pulse every 3 min. During post-processing, we analyzed animal activity around each stimulation timepoint. For the 10 s prior to each stimulation timepoint, we classified animals as quiescent, slow-moving, or fast-moving. Two categories for wake behavior (i.e. slow-moving and fast-moving) were necessary because a behavioral response could not be detected in animals that were already moving with a high activity. That is, an animal that was clearly awake and moving with high activity in the microfluidic chamber did not display increased activity after receiving mechanical stimuli. However, a detectable behavioral response was apparent in wake, but low-activity animals. We used an activity threshold of < 0.08 for quiescent animals, 0.08-0.35 for low-activity animals, and > 0.35 for high-activity animals (the average animal activity across all traces was 0.36). This classification was performed for each stimulation (20 stimulations for each animal). These classified groups comprise the heatmaps showed in Figure 2B. Note that because animals are more likely to be in the wake state, this classification method leads to lower n-values for the sleep condition. From each animal in each group, we also calculated the mean activity for 10 s post-stimulation, excluding a 2 s period where the microfluidic valves caused movement artifacts, which make up the data shown in Figure 2C.

### Homeostasis/RIS Optogenetic Inhibition

This protocol was adapted from Wu *et al* (2018). HBR1374 animals expressed the light-gated proton pump ArchT in the RIS neuron and were raised on NGM seeded with 0.2 mM all-trans retinal (+ATR). Control animals were also the HBR1374 strain but were not raised in ATR (- ATR). Experiments were performed on an inverted Nikon microscope. Animals were loaded in 110 μm-wide chambers (see Figure 6 inset) and imaged at 4 fps with a Basler Ace (acA1920-40um) CMOS camera. Experiments lasted 2 hr. Between 1-1.5 hr, we illuminated animals with 565 nm light (3 mW/mm^2^).

### Whole-Brain Imaging in Behaving Animals

Microfluidic devices were fabricated as previously described (see “Microfluidic Device Fabrication” subsection), and geometries were modeled off of previous methods for whole-brain imaging ^32, 33, 65, 69, 70^; however, animals were not chemically paralyzed. We used transgenic animals with pan-neuronal expression of GCaMP6s in the nuclei of all neurons ^66, 68^. Experiments were performed on an inverted Nikon microscope with a 40X water-immersion objective (NA = 1.15). An Andor Zyla 4.2 USB 3.0 sCMOS captured the images at 5 fps (50 ms exposures, 2x2 binning). Because blue light can wake *C. elegans*, when animals were immobilized in the chambers, we first initialized a “habituation” protocol. Here, we flashed the 460 nm excitation light for 0.15 s at 1 Hz for 30 min. This habituated animals without photobleaching the calcium indicator. After the habituation period, we imaged continuously for 15 min, but only analyzed the first 10 min of data due to photobleaching. With this protocol, we captured a sleep-wake transition in ∼25% of animals.

To analyze data, we chose 10 neurons at random to track by hand in FIJI. An 8x8 pixel ROI was then used for each neuron in each frame. A neuron near the center of the ganglia was used to draw an ROI around the head ganglia to calculate average brain fluorescence in each frame. We calculated animal activity in two ways: by frame-by-frame subtraction (sampling the frames a 3 Hz), and by calculating the average displacement of the tracked neurons. Averaged and normalized neuron displacement make up the behavioral traces in Figure 4, **Supplementary Movie 6**. Both methods were prone to fluctuating baselines that were not present during behavioral-only recordings, making it difficult to set a standard threshold across animals.

However, clear quiescent periods were subjectively apparent. Therefore, for each animal we plotted only the animal behavioral activity and selected sleep bouts manually. As before, only bouts longer than 30 s were counted. We then calculated the mean ganglia and neuron fluorescence during sleep and wake for each worm.

Behavioral activity was smoothed over 3 s and normalized as previously described (see “Behavioral Quantification and Sleep Detection” subsection). Raw ganglia and neuron activity was smoothed over 3 s before calculating ΔF/F. We denote ΔF/F as [F(t) – F0(t)]/F0(t) where F0(t) is the minimum fluorescence value up to time *t*.

### Whole-Brain Imaging in Paralyzed Animals

Whole-brain imaging in paralyzed animals closely followed the protocol for behaving animals (see “Whole-Brain Imaging in Behaving Animals” subsection), however 5 mM tetramisole was added to the buffer to chemically paralyze animals, similar to previously reported methods ^32, 33, 65, 69^. In addition, the 40X objective was mounted onto a piezo scanner (Physik Instrumente). We used MATLAB to control a data-acquisition box (National Instruments) that output voltage waveforms to trigger both the piezo scanner and Zyla camera. The DAQ output a sawtooth waveform to the piezo, which enabled scanning across 30 μm in the z-axis. At 12 different points along the scan, the piezo would briefly settle for 30 ms while the DAQ also triggered a 10 ms camera frame acquisition. This approach led to a volumetric imaging rate of 2.84 vol/s over 10 min periods.

Following imaging, we used a custom MATLAB GUI to manually choose neuron locations. The GUI allowed users to look at max projections, volumes, or single imaging planes across the duration of the recording. Consecutive planes or volumes could also be averaged to reduce noise. This allowed for the x-y-z positions of neurons to be manually located and stored. The average of a 3x3x3 pixel ROI was used around each neuron when extracting mean fluorescence. Raw neuron activity was smoothed over 3 s before calculating ΔF/F. We denote ΔF/F as [F(t) – F0(t)]/F0(t) where F0(t) is the minimum fluorescence value up to time *t*.

### RIS Imaging

7-10 animals per trial (6 trials total) were confined and imaged simultaneously under 10X magnification in the 50 μm-wide chamber geometry. HRB1361 animals expressed both GCaMP3 and mKate in RIS neurons under the FLP-11 promoter. Experiments took place on an inverted Nikon microscope, with dual excitation and emission for simultaneous two-color imaging (FF01-468/553-25 and F01-512/630-25 BrightLine dual-band bandpass filters for excitation and emission, respectively). An X-Cite XLED1 light source provided both 460 nm and 565 nm excitation light and a Tucam image splitter (Andor) split the mKate and GCaMP channels onto two Andor Zyla 4.2 CMOS cameras. We imaged animals for 2 hr and used a data-acquisition box (National Instruments) controlled with custom MATLAB scripts to simultaneously trigger 10 ms camera exposures and 30 ms XLED flashes at 0.5 Hz (3x3 binning). Following experiments, we flushed animals from the device and recorded the background for each color channel.

For post-processing, we used the mKate channel to quantify behavior and track the location of RIS neurons. After subtracting background, we calculated animal activity by drawing an ROI around each animal and performing the same frame-by-frame subtraction method as previously described (see “Behavioral Quantification and Sleep Detection” subsection), but here we used a with a pixel change threshold of 400 because of the high dynamic range of the Zyla sensor. After normalizing behavioral activity, a threshold of 0.2 was used to detect sleep bouts, with a minimum sleep time of 30 s. To detect the RIS location in each frame, we again subtracted the image background. We then isolated and binarized each animal ROI, leaving only the RIS neuron and, occasionally, pieces of its processes. The largest object in the ROI was the RIS soma, which we used to attain the soma centroid. To get the average RIS fluorescence in the frame, we drew a 25x25 pixel ROI around the centroid in each channel and averaged the 20 largest pixel values. We normalized this raw fluorescence in each channel by calculating ΔF/F = [F(t) – F0(t)]/F0(t), where F0(t) is the minimum fluorescence value up to time *t*. Following this normalization, we calculated the ratio of each color channel R = (ΔF/F)GCaMP/(ΔF/F)mKate. Finally, the reported ratio values are ΔR/R = [R(t) – Ro]/Ro, where Ro is the lower 20^th^ percentile value.

Using behavioral data (from frame-by-frame subtraction), we detected sleep bouts and calculated the average RIS ΔR/R activity during wake and sleep (Figure 5C). To calculate correlation values, across all animals we split behavioral and fluorescence data into 10 min windows and calculated the fluorescence correlation with behavior in each window. ΔR/R data showed a strong negative correlation with behavior (Figure 5D, green trace). Randomly shuffling the ΔR/R windows as a control removes this strong correlation (Figure 5D, grey trace). To test for movement artifacts that could possibly lead to a correlation between fluorescence and behavior, we performed the same unshuffled analysis with the mKate channel only (Figure 5D, red trace).

### Environmental Control

The microfluidic setup during environmental manipulation was identical the previously stated (see “Standard Microfluidic Behavioral Assays” subsection); however, for each condition we manipulated individual aspects of the environment. The baseline phenotype in 500 μm-wide chambers (**Supplementary Movie 1**, Figure 6) was chosen because animals could move freely in a manner similar to swimming, thereby reducing the overall effect of the microfluidic confinement.

Most assays involved transferring animals directly from seeded NGM into the microfluidic device. However, under “Starved” conditions we used a hair pick to remove animals from seeded NGM, washed them in a droplet of M9 buffer, then placed them onto a fresh, unseeded NGM plate. After 2 hr, animals were loaded into the microfluidic device with no food in the buffer. Therefore, animals were deprived of a food source for 2 hr when imaging began and did not have access to food during the experiment.

Most assays were also performed with no food in the M9 buffer. However, under “+Food” conditions, we dissolved 4.56 mg/mL of freeze-dried OP50 (LabTie) into the M9 buffer, providing a food source for animals throughout the imaging time.

In “+Compression” conditions, animals were confined to 50 x 1100 um-wide microfluidic chambers (**Supplementary Movie 2**).

As previously stated, the standard imaging temperature due to LED lighting was 21-22 °C (see “Standard Microfluidic Behavioral Assays” subsection). We controlled temperature in the same manner as heat-shock experiments, using Peltier devices to heat or cool the microfluidic chips to 24-25 °C or 17-18 °C, respectively.

### DAF-16::GFP Imaging

Experiments took place over the course of 2 hr on Nikon inverted scope with an Andor Zyla 4.2 USB 3.0 sCMOS and 4X magnification for multi-worm imaging. Environmental control took place as described in the “Environmental Control” subsection. Every 5 min, we captured 7 frames (20 ms exposures) at 1 Hz with 1x1 binning. Otherwise, animals were in darkness. Following imaging, we flushed animals from the chamber and recorded background frames.

Puncta quantification took place similar to previous methods with custom MATLAB scripts ^80^. We subtracted the background from each frame and smoothed the image with a median filter. We thresholded this image to detect animal bodies and drew a ROI around each. The smoothed image was further processed with a 3x3 high frequency filter, which enhanced the contrast of puncta and allowed us to binarize the image to detect most DAF-16 aggregations in the frame. For each animal ROI, we then found puncta by counting all objects with a size between 2 and 30 pixels. To mitigate the effects of animal movement, which caused some animals to appear blurred and reduced puncta count, of the 7 frames that were captured every 5 min, we averaged the 3 largest puncta counts for each ROI. This average makes up the final puncta count for each animal at the given timepoint. The puncta count reported (**Supplementary Figure 6**) are likely lower than the true count that would be attained with high-magnification imaging. Low-magnification not only increases the number of animals imaged per trial but was also necessary to capture the full length of the large microfluidic chambers. Because we applied the same algorithm to all environmental conditions, we expect the reported trends to remain true in high-magnification conditions.

### Quantification and Statistical Analysis

Boxplots with scatterplots overlaid were constructed in MATLAB. For the boxplots, the top and bottom edges of the box indicated the 25^th^ and 75^th^ percentiles, respectively; the whiskers extend to the most extreme data points not calculated as outliers; and the grey line in the box indicates the median.

All statistical tests are indicated in figure legends. Almost all statistical comparisons were between more than 2 groups with unequal variances. Thus, the non-parametric Kruskal-Wallis test with a *post-hoc* Dunn-Sidak test was used to test for statistical significance. For data in which we tested whether one group displayed phenotypic differences (either behavioral or neural) during wake and sleep, we used the paired t-test.

## DATA AVAILABILITY

All raw data (i.e. raw animal activities from frame-by-frame subtraction) will be made publicly available at [insert BOX link upon acceptance]. Raw video files and code for analysis will be provided upon request.

## SUPPLEMENTAL INFORMATION

**Supplementary Figure 1.**
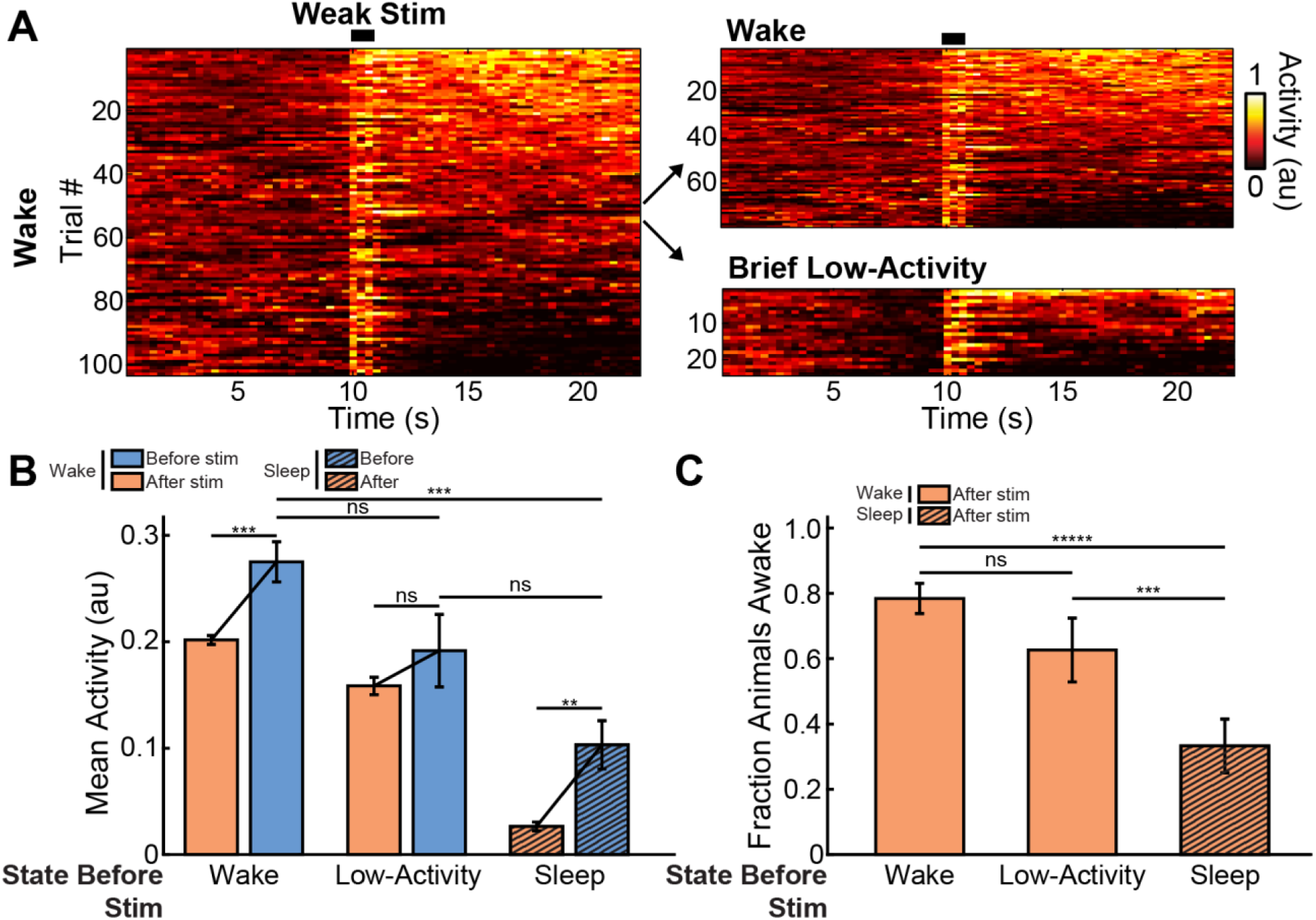
Further comparison of behavioral responses to weak mechanical stimulation. *(A) (Left)* Heatmap of all behavioral activity from all wake animals receiving a weak mechanical stimulus (n = 176 total trials, note that in Figure 2 only the first 25 animals are shown for clarity). *(Right)* We further parsed this data set into “Wake” and “Low-Activity” states, where “Low-Activity” is defined as having an average behavioral activity level below the sleep threshold for a short 3 s period immediately before the stimulus. *(B)* Average behavioral activity before and after the weak stimulus for animals in the Wake and Low-Activity states prior to the stimulus. We also compared these results to animals in the Sleep state prior to the stimulus (same data as Figure 2D, error bars are sem, ***p < 0.001, Kruskal-Wallis with a *post-hoc* Dunn-Sidak test). *(C)* Fraction of animals awake following the stimulus for the Wake and Low-Activity states. We also compared these data to the Sleep state (same data as Figure 2E). Animals in the Sleep state are less likely to respond to weak stimuli and transition to wakefulness, compared to animals in the wake state regardless of their activity level prior to the stimulus. (Error bars are standard deviation, calculated by bootstrapping each data set with 5000 iterations; ns = not significant, ***p<0.001, ****p<0.0001; significance was calculated by data resampling 5000 iterations and a *post hoc* Bonferroni correction).

**Supplementary Figure 2.**
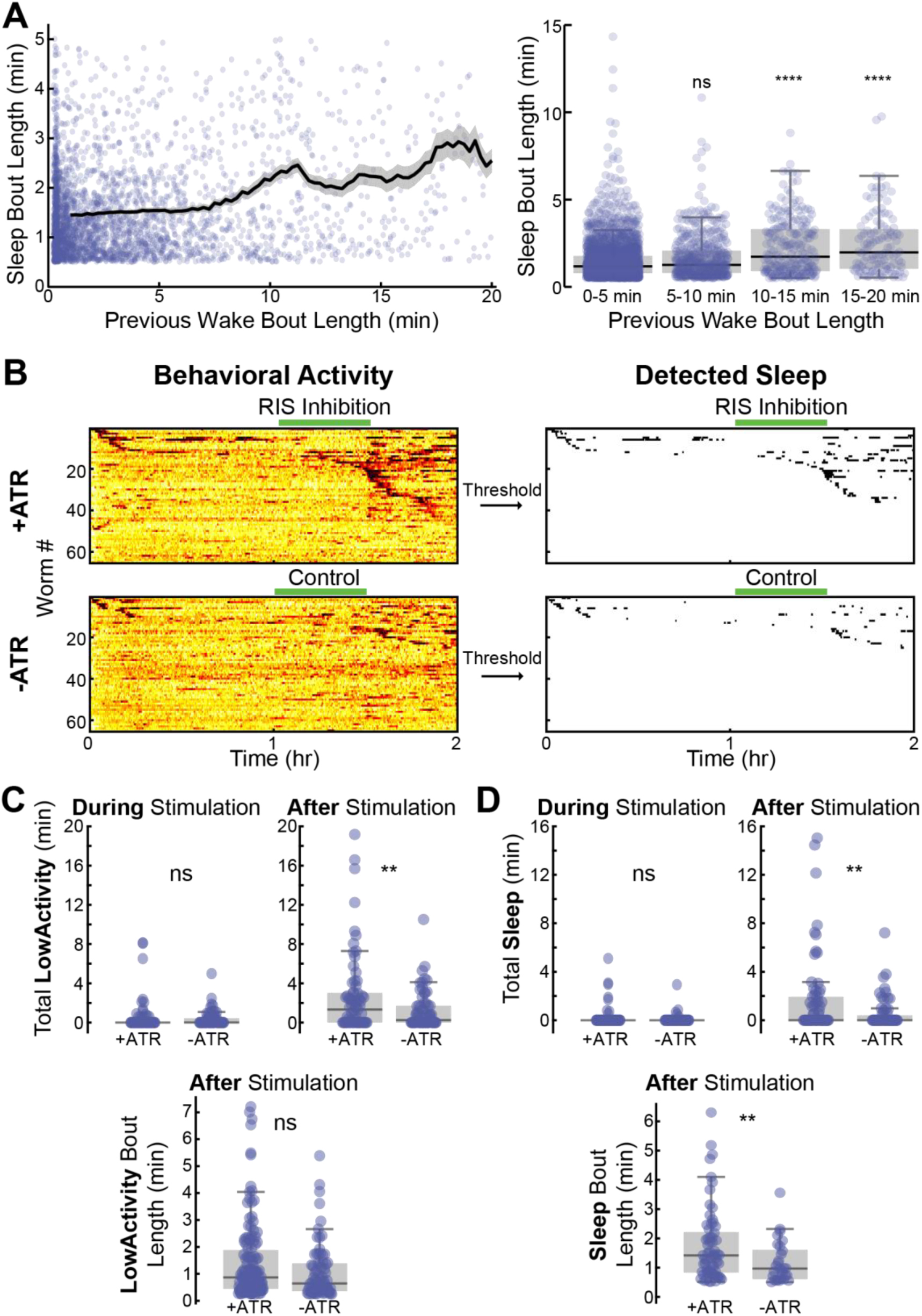
Evidence for homeostasis in microfluidic-induced sleep. *(A)* Sleep bout length is dependent on the length of the previous wake bout. *(Left)* Data points represent individual sleep and wake bout pairs from data collected from 50 and 500 μm chambers with WT animals (see Figure 1, 4 hr-long recordings and Figure 6, 2 hr-long recordings). *(Right)* Wake bout lengths binned into 5 min windows. As the wake bout length increases, the following sleep bouts increase in length, suggesting a micro-homeostatic mechanism (ns = not significant, ****p<0.0001 compared to 0-5 min data set, Kruskal-Wallis with a *post-hoc* Dunn-Sidak test). *(B-D)* Animals showed a homeostatic rebound in response to prolonged RIS optogenetic inhibition. *(B) (Left)* Heatmaps of animal behavioral activity during an optogenetic assay where green light illumination was delivered between 1-1.5 hr. *(Right)* Raster plots of detected sleep bouts. +ATR animals were grown with OP50 seeded with *all-trans retinal*. -ATR animals were not, thus making optogenetic inhibition ineffective. *(C)* We used a threshold 66% higher than the sleep threshold to detect all low-activity bouts (these include sleep and other low-activity behaviors). +ATR animals showed similar behavioral activity during the stimulus, but more low-activity after the stimulus, suggesting that prolonged RIS inhibition led to a homeostatic response following optogenetic inhibition of the sleep-promoting RIS neuron. *(D)* Using our standard threshold for sleep detection, +ATR and -ATR animals surprisingly showed no significant difference in total sleep during illumination. However, +ATR animals showed significantly more sleep following stimulation, suggesting a homeostatic response to prolonged RIS inhibition. In addition, +ATR animals showed longer sleep bouts following illumination. These results suggest that *C. elegans* show increased microfluidic-induced sleep and low-activity following extended RIS inhibition. (ns = not significant, *p<0.05, **p<0.01, unpaired two-sided t-test).

**Supplementary Figure 3.**
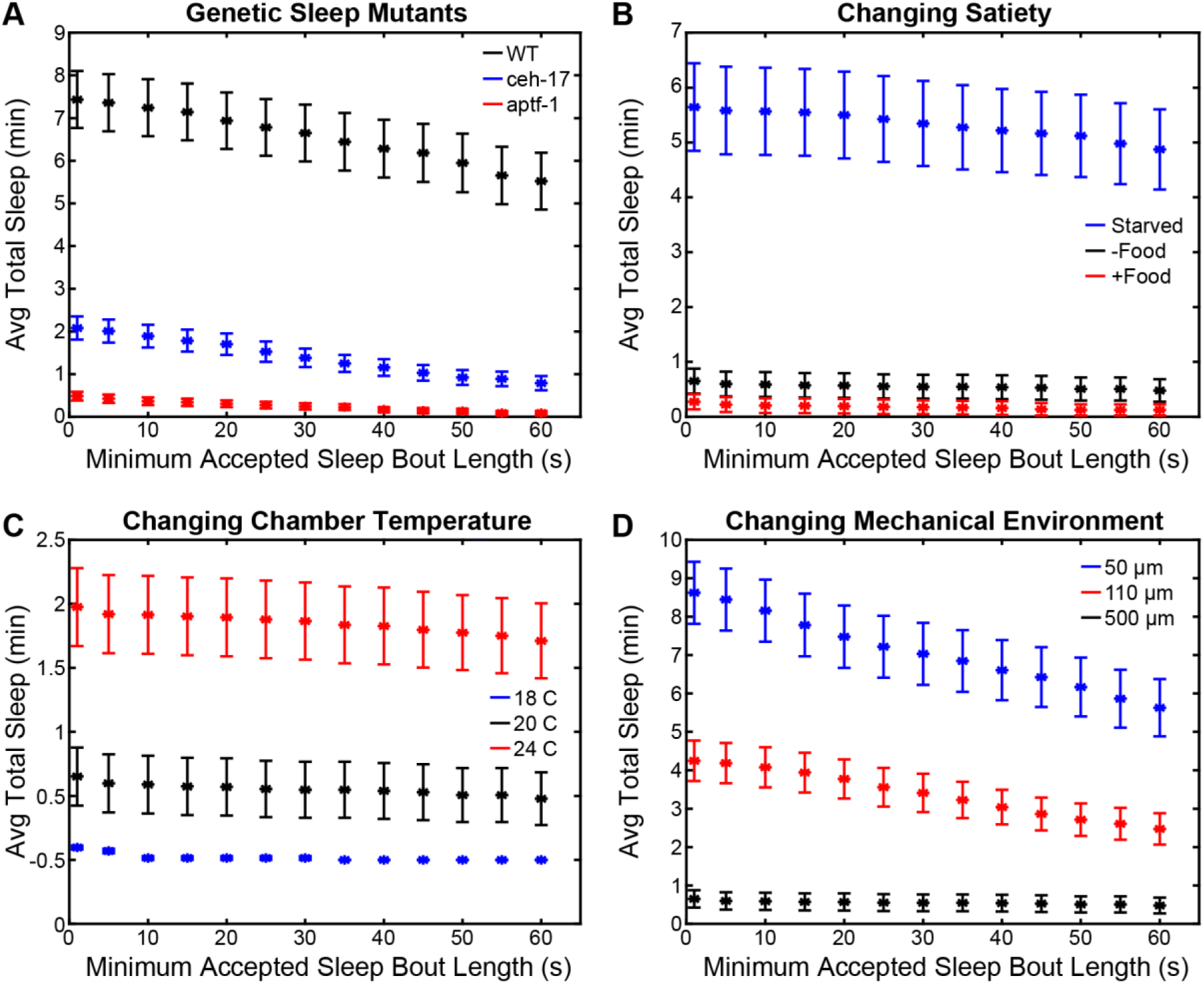
Reported sleep trends are unaltered by the choice of minimum sleep time. For all data reported, low-activity bouts lasting less than 30 s were removed in order to reduce false-positive detections (see Methods). Here, we show that reducing this value to even 1 s does not change the reported conclusions. Data sets analyzed here are from various portions of the manuscript. *(A)* Genetic mutant data reported in Figure 2 (chamber widths are 50 μm). *(B)* Satiety data from Figure 6 (chamber widths are 500 μm). *(C)* Temperature data for WT animals in Figure 6 (chamber widths are 500 μm). (*D)* Data from WT animals in which chamber width was changed in Figure 6 (chamber widths vary and are indicated in the legend). As expected, data from large chambers (500 μm width) is highly stable due an easy distinguishing between sleep and wakefulness. More variation is seen in smaller chambers (50 μm width), where animal movement is heavily restricted, and animals exhibit more low-activity bouts that are not necessarily sleep behavior. In all cases however, the reported trends in throughout the manuscript are sound.

**Supplementary Figure 4.**
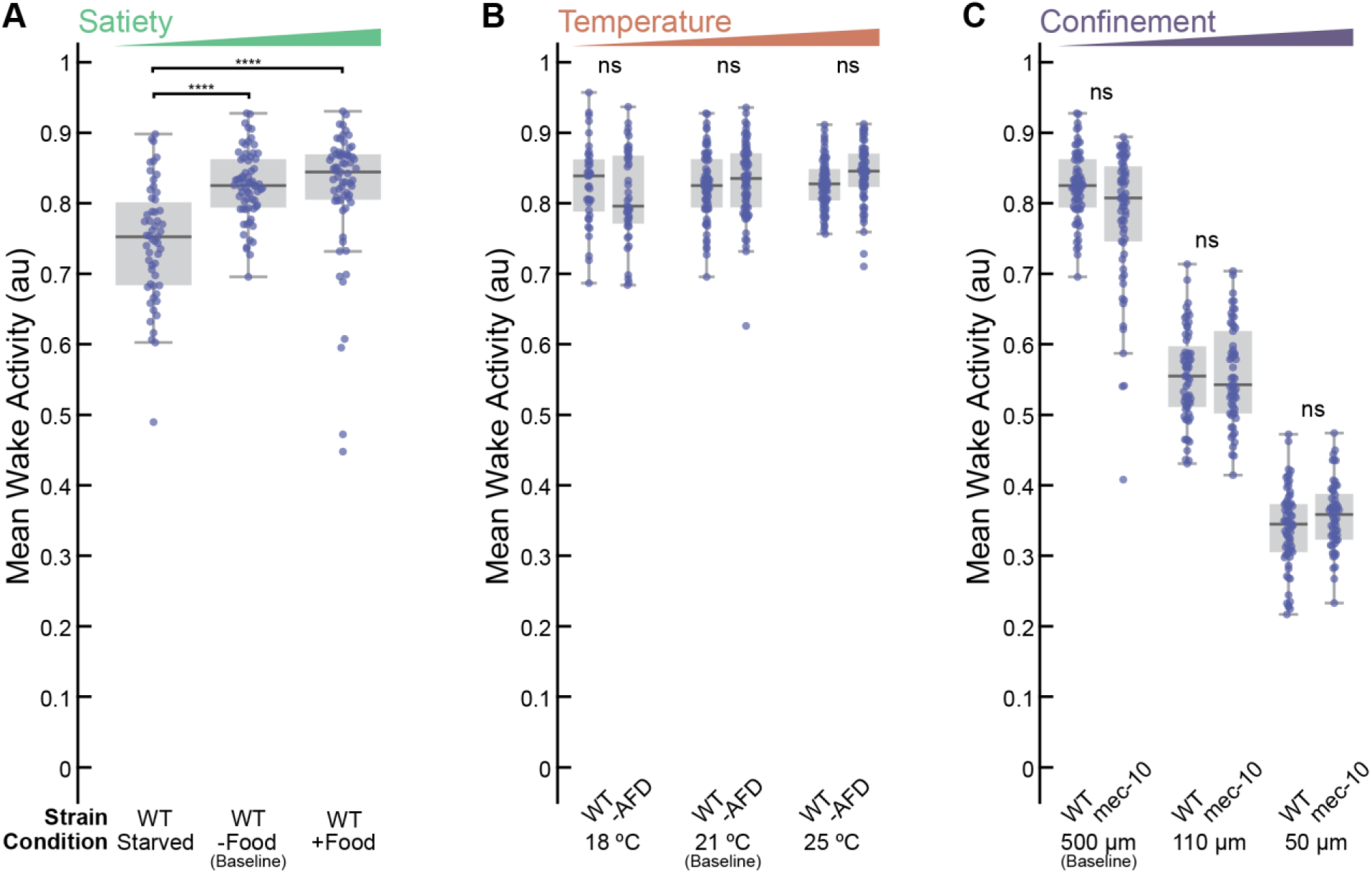
Behavioral activity during wakefulness in different environmental conditions. Data is the averaged wake-state behavioral activity for each animal in Figure 6. *(A)* Only WT animals in the starved condition only show less activity compared to -Food and +Food. *(B)* No significant differences were found in either WT or *gcy-23(nj37);gcy-8(oy44);gcy-18(nj38)* mutants (labeled as -AFD) at any microfluidic device temperature. *(C)* As expected, restricting animal movement significantly changes behavioral activity during wakefulness. However, no changes were observed between WT and *mec-10(tm1552)* animals. (****p<0.0001, ns = not significant, Kruskal-Wallis with a *post-hoc* Dunn-Sidak test).

**Supplementary Figure 5.**
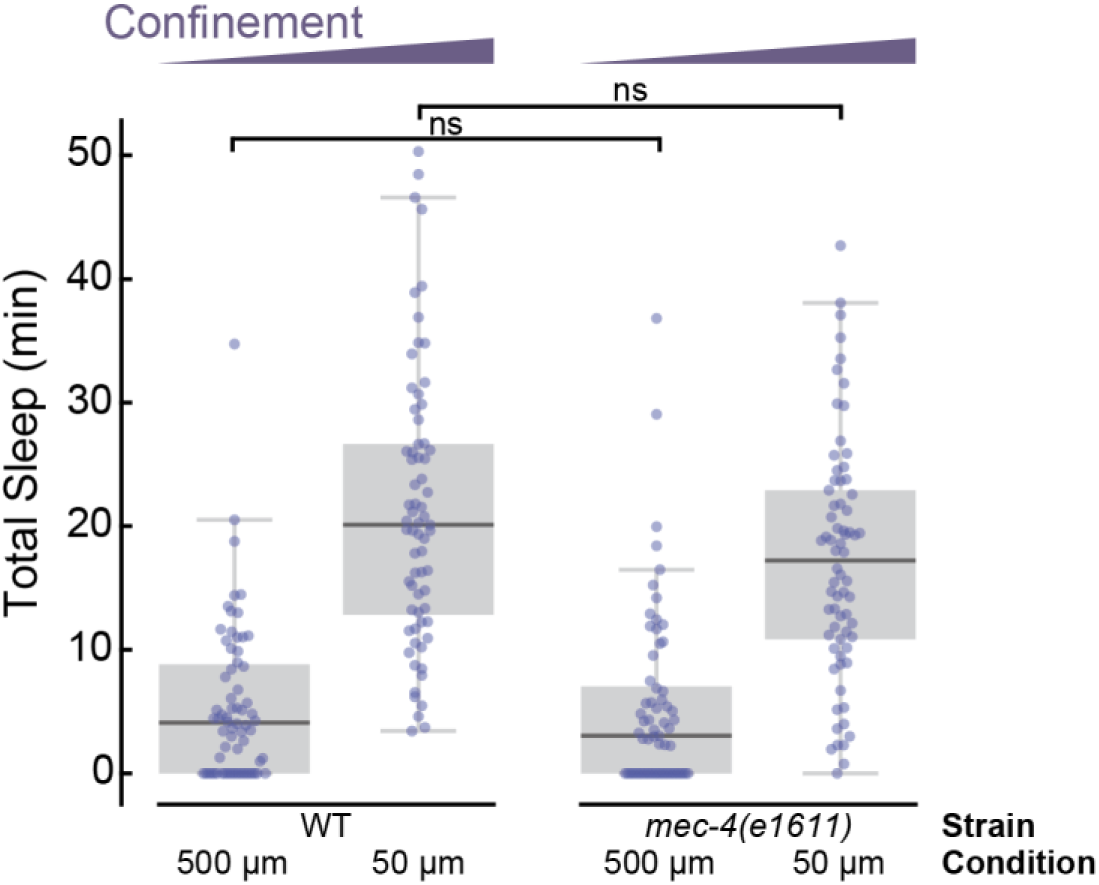
*mec-4(e1611)* gentle-touch-defective mutants show same confinement phenotype as WT. When animals are allowed to swim in large 500 μm-wide chambers, mutant and WT strains show the same amount of total sleep. When partially immobilized in 50 μm-wide chambers, each strain shows an increase in total microfluidic-induced sleep, but the phenotypes remain the same. These results indicate that harsh-touch mechanosensory pathways, such as *mec- 10(tm1552)* (Figure 6), drive microfluidic-induced sleep. (WT 500 μm n = 68, WT 50 μm n = 69, *mec-4(e1611)* 500 μm n = 65, *mec-4(e1611)* 500 μm n = 70; ns = not significant, Kruskal-Wallis with a *post-hoc* Dunn-Sidak test).

**Supplementary Figure 6.**
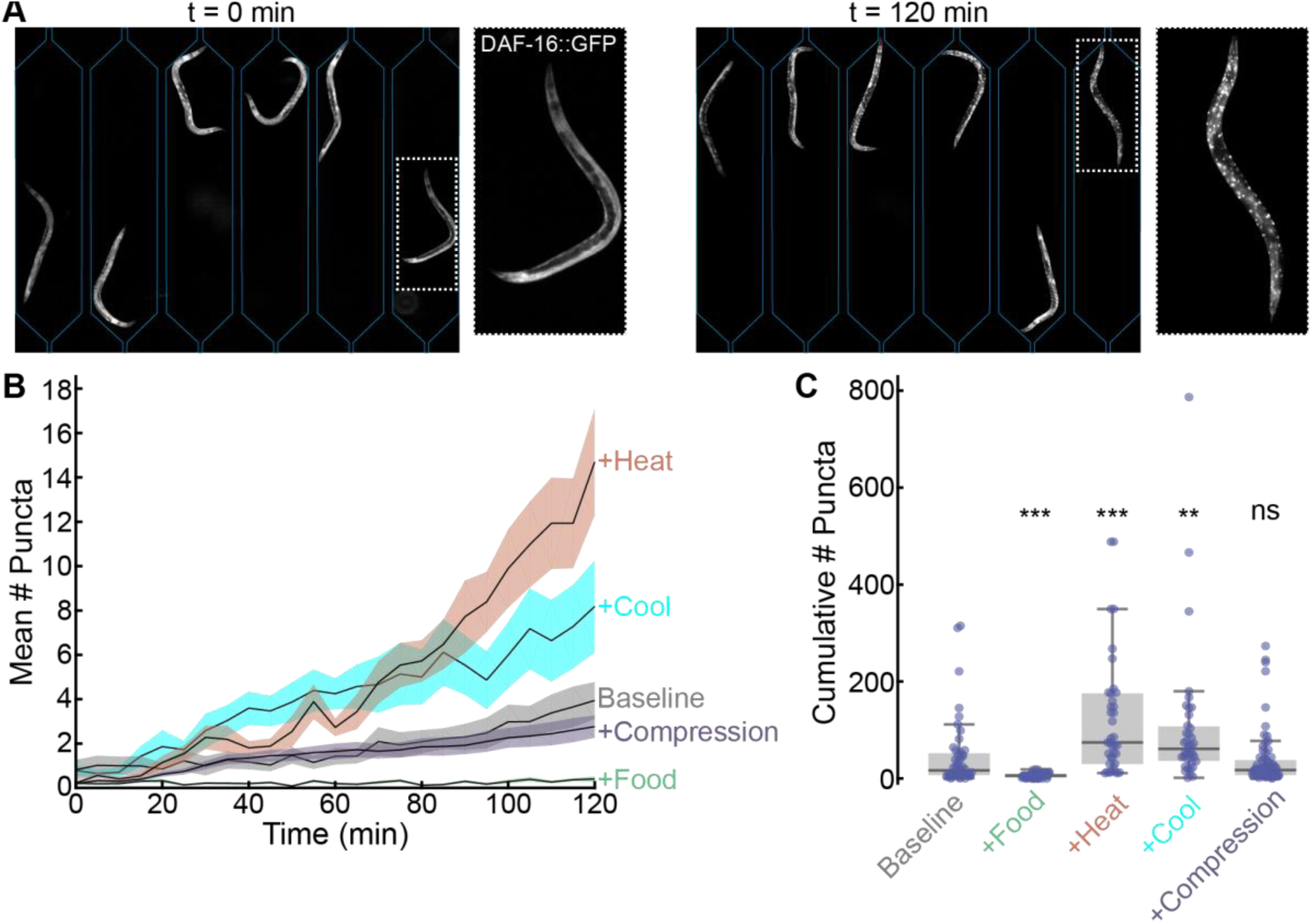
DAF-16::GFP imaging shows that microfluidic-induced sleep partially correlates with *C. elegans* stress. *(A)* Fluorescent micrographs (with background subtracted) of DAF16::GFP animals confined to 500 μm-wide chambers during the +Heat condition. Microfluidic chambers are outlined. Left image (t = 0 min) shows diffuse DAF-16::GFP. Over the course of 2 hr, DAF-16 localizes to the nucleus (right image). *(B)* Puncta formation with respect to time for Baseline, +Food, +Heat, +Cool, and +Compression conditions (see Methods, Figure 6). *(C)* Cumulative number of puncta during imaging. Individual data points represent individual animals. (Baseline n = 48, +Food n = 46, +Heat n = 38, +Cool n = 39, +Compression n = 71; **p < 0.01, ***p < 0.001, ns = not significant, Kruskal-Wallis with a *post-hoc* Dunn-Sidak test).

**Supplementary Figure 7.**
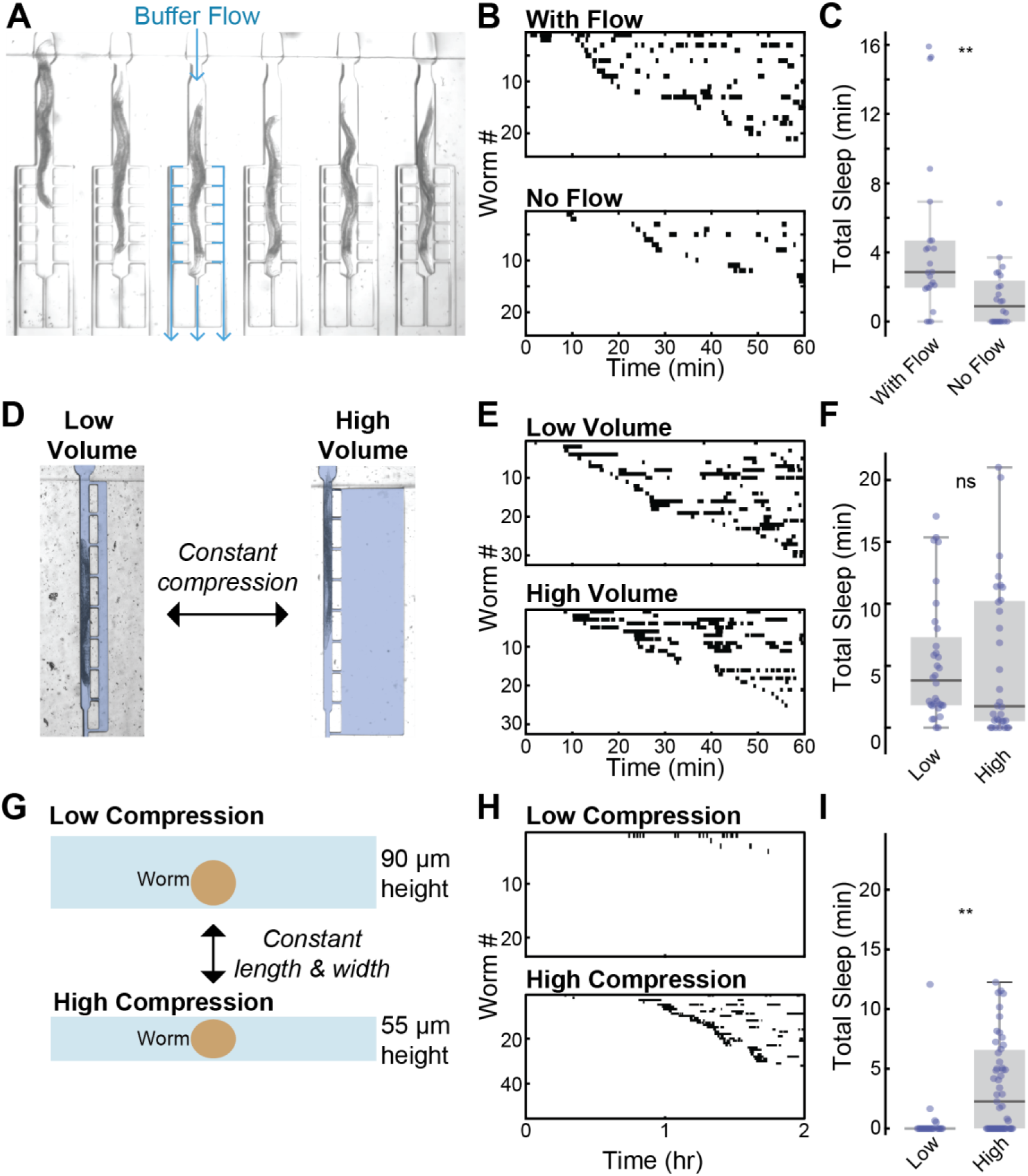
Waste buildup, chamber volume, and stress from loading do not strongly influence microfluidic-induced sleep. *(A)* Image of animals confined in microfluidic chambers designed for constantly flowing buffer to stabilize O2 concentrations and remove the buildup of CO2 and other byproducts. Blue paths indicate the direction of flow for a single chamber. Flow rate was ∼1 mL/hr. *(B)* Raster plots of detected sleep with and without flow using the geometry in (A). *(C)* We surprisingly observed more sleep with the buffer flow, indicating microfluidic-induced sleep is likely not driven by changing gas concentration levels biological byproducts (n = 24 for each condition). *(D)* Chamber designs for maintaining a constant animal compression while changing fluidic volume. Fluid is false-colored in pale blue. *(E)* Raster plots of detected sleep for each chamber type. *(F)* Animals in chambers of different volume do not show different amounts of sleep (n = 32 for each condition). *(G)* Schematic of chamber cross section that have same width and length, but different chamber heights. *(H)* Raster plots of detected sleep. *(I)* Sleep in the 90um tall chambers is essentially abolished, demonstrating that the animal loading process and microfluidic environment alone do not drive sleep (n = 22). Sleep dramatically increases in the 55 um tall chambers (n = 50), again demonstrating that the mechanical environment regulates sleep strongly. (ns = not significant, **p < 0.01, unpaired two-sided t-test).

**Supplementary Figure 8.**
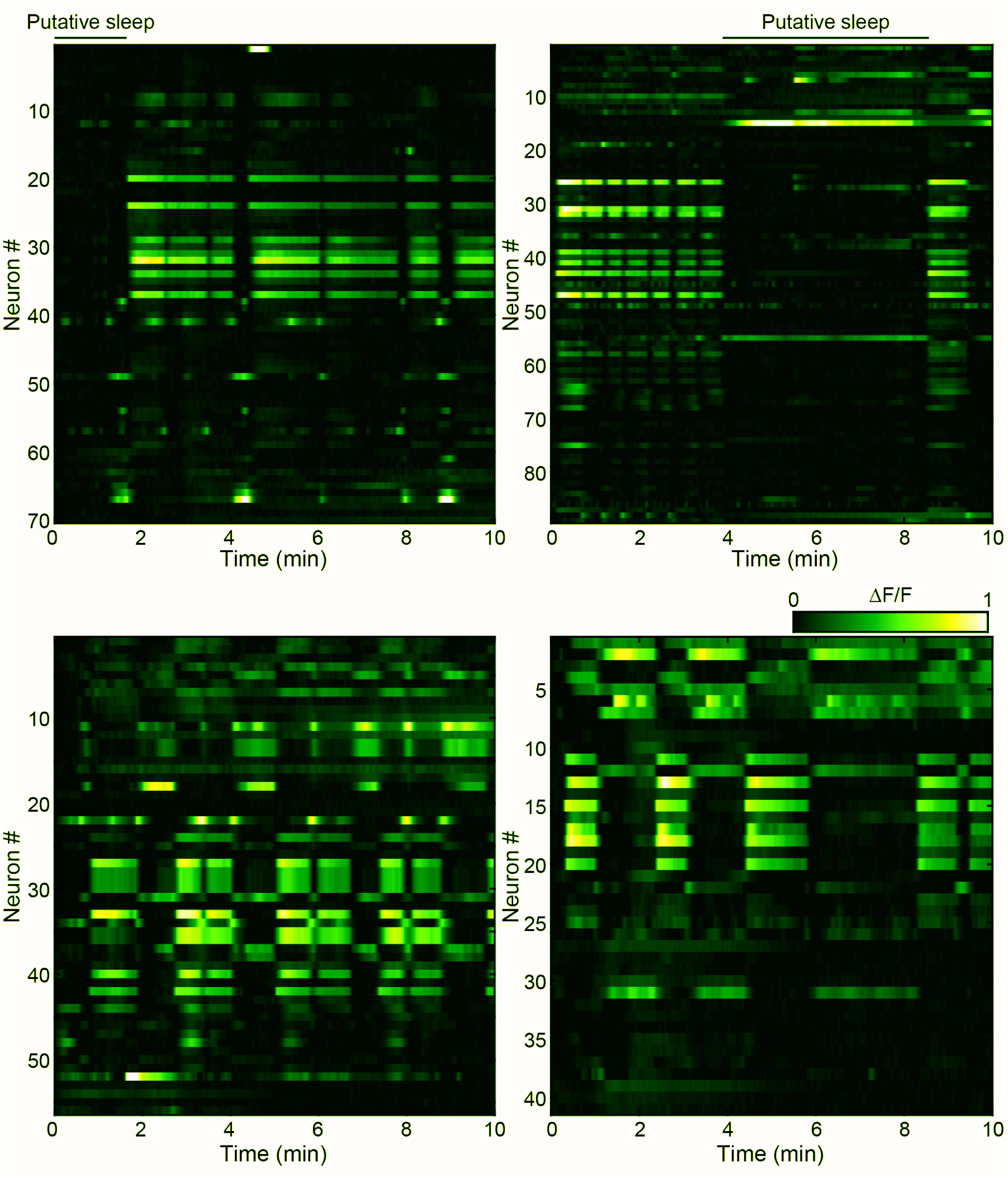
Whole-brain imaging in paralyzed animals. Representative volumetric imaging data from four paralyzed animals during whole-brain imaging. The majority of the data resembles previous imaging work during wakefulness ^32, 33^, wherein many neurons show correlated calcium dynamics. Putative sleep states (top two panels) are labeled but cannot be confirmed without behavioral readouts. These data demonstrate that microfluidic-induced sleep can be used as a model behavior for understanding how brain-wide neural circuits drive spontaneous brain state transitions.

**Supplementary Movie 1.** microfluidic-induced sleep in a large microfluidic chamber. (Top) Animal activity calculated by subtracting consecutive frames (see Methods). (Bottom) Video (with background subtracted) of the animal swimming in large microfluidic chamber. The animal is initially in the active state, then spontaneously transitions in and out of quiescence.

**Supplementary Movie 2.** microfluidic-induced sleep in a small microfluidic chamber. *(Top)* Animal activity trace calculated by subtracting consecutive frames (see Methods). *(Bottom)* Video (with background subtracted) of the animal partially immobilized in a small microfluidic chamber. The animal is initially in the active state, then spontaneously transitions in and out of quiescence.

**Supplementary Movie 3.** microfluidic-induced sleep is reversible with strong blue light illumination. *(Top)* Animal activity trace. Blue bar indicates a 5 s light stimulation. *(Bottom)* Video (with background subtracted) of an animal partially immobilized in a small microfluidic chamber. The animal is initially quiescent but wakes upon illumination.

**Supplementary Movie 4.** microfluidic-induced sleep is reversible with strong mechanical stimulation. Two animals receive strong mechanical stimulation via a microfluidic push-down valve. The first animal is in the wake state, but behaviorally responds to stimulation. The second animal is in the sleep state and wakes upon stimulation.

**Supplementary Movie 5.** Animals show a decreased response to weak mechanical stimuli during microfluidic-induced sleep. Two animals receive weak mechanical stimulation via a microfluidic push-down valve. The first animal is in the wake state, but still behaviorally responds to stimulation. The second animal is in the sleep state. Due to an increased arousal threshold, the animal does not respond.

**Supplementary Movie 6.** Two representative examples of a brain and behavioral state transitions during microfluidic-induced sleep. Video shows single-plane, whole-brain epifluorescence of GCaMP6s imaging during a microfluidic-induced sleep transition. Top trace is animal behavioral activity. Middle trace shows the average ganglia fluorescence. Bottom trace shows the fluorescence of ten individual neurons.

## Notes

#### Summary of Updates

New analyses added to Figure 2. Several new supplemental experiments and analyses. New author added.

